# *Drosophila* axis extension is robust to an orthogonal pull by invaginating mesoderm

**DOI:** 10.1101/2023.07.18.549479

**Authors:** Claire M Lye, Guy B. Blanchard, Jenny Evans, Alexander Nestor-Bergmann, Bénédicte Sanson

## Abstract

As tissues grow and change shape during animal development, they physically pull and push on each other and these mechanical interactions can be important for morphogenesis. During *Drosophila* gastrulation, mesoderm invagination temporally overlaps with the extension of the ectodermal germband; the latter is caused primarily by Myosin II-driven polarised cell intercalation. Here we investigate the impact of mesoderm invagination on ectoderm extension, examining possible mechanical and mechanotransductive effects on Myosin II recruitment and polarised cell intercalation. We find that the germband ectoderm is deformed by the mesoderm pulling in the orthogonal direction, showing mechanical coupling between these tissues. However, we do not find a significant change in Myosin II planar polarisation in response to mesoderm invagination, nor an effect on the rate of junction shrinkage leading to cell intercalation events. We find some impact on the orientation of neighbour exchange events, and an increased rate of growth of new cell junctions, but this makes little difference to the rate of cell intercalation. We conclude that the cellular mechanisms of axis extension are robust to the mechanical pull of mesoderm invagination.

## INTRODUCTION

The generation of tissue shapes during animal development is complex, but we are beginning to understand the fundamental cell behaviours underlying this process such as oriented cell division, cell shape changes and changes in relative cell positions. Many of these processes can be controlled cell autonomously, by forces generated within the cell, but it is becoming increasingly clear that they can also be influenced by extrinsic forces generated by the movements of neighbouring tissues. For example, extrinsic forces have been implicated in driving cell shape changes, cell intercalation and the reorganisation of planar polarity in developing tissues (Butler et al., 2009; Lye et al., 2015; Caussinus et al., 2008; Blackie et al., 2021; Aigouy et al., 2010; Aw et al., 2016; Collinet et al., 2015). In addition to examples where physical interactions between tissues are important for their morphogenesis, it is becoming apparent that mechanisms also exist to buffer physical forces and mechanically isolate tissues from one another (Rauzi et al., 2015; Duda et al., 2019; Ashour et al., 2023). Despite this interest in the physical interactions between tissues, we have yet to gain a thorough understanding of to what extent forces are transmitted across tissues, and how this is regulated to allow embryos to undergo robust and reproducible morphogenesis.

Early embryogenesis in *Drosophila* has become an important paradigm for understanding how tissue morphogenesis is driven by a combination of forces generated directly by cells (intrinsic forces) and indirectly, at the tissue or embryo scale (extrinsic forces), as several well-characterised morphogenetic movements occur within a short period of time (Gustafson et al., 2022; Fierling et al., 2022; Dicko et al., 2017; Rauzi et al., 2015; Lye and Sanson, 2011; Martin et al., 2010). Specifically, at the same time as the mesoderm invaginates ventrally and the endoderm invaginates posteriorly, the ectoderm initiates convergence and extension to extend the main body axis of the embryo (germband extension, GBE) (Fig. 1A-E) (Lye et al., 2015; Gheisari et al., 2020). GBE is caused primarily by polarised cell intercalation driven by Myosin II (Bertet et al., 2004; Zallen and Wieschaus, 2004; Collinet and Lecuit, 2021). Antero-posterior (AP) patterning controls the planar polarised distribution of junctional cortical Myosin II, which leads to shrinkage of cell junctions parallel to the dorso-ventral (DV) axis (so called DV-oriented junctions) leading to cells exchanging neighbours (Irvine and Wieschaus, 1994; Bertet et al., 2004; Zallen and Wieschaus, 2004). In addition to this primary mechanism, an extrinsic AP-oriented pull from the invaginating posterior endoderm also contributes to axis extension, by elongating cells and also orienting growing junctions during polarised cell intercalation (Collinet et al., 2015; Butler et al., 2009; Lye et al., 2015; Kong et al., 2017).

**Figure 1:**
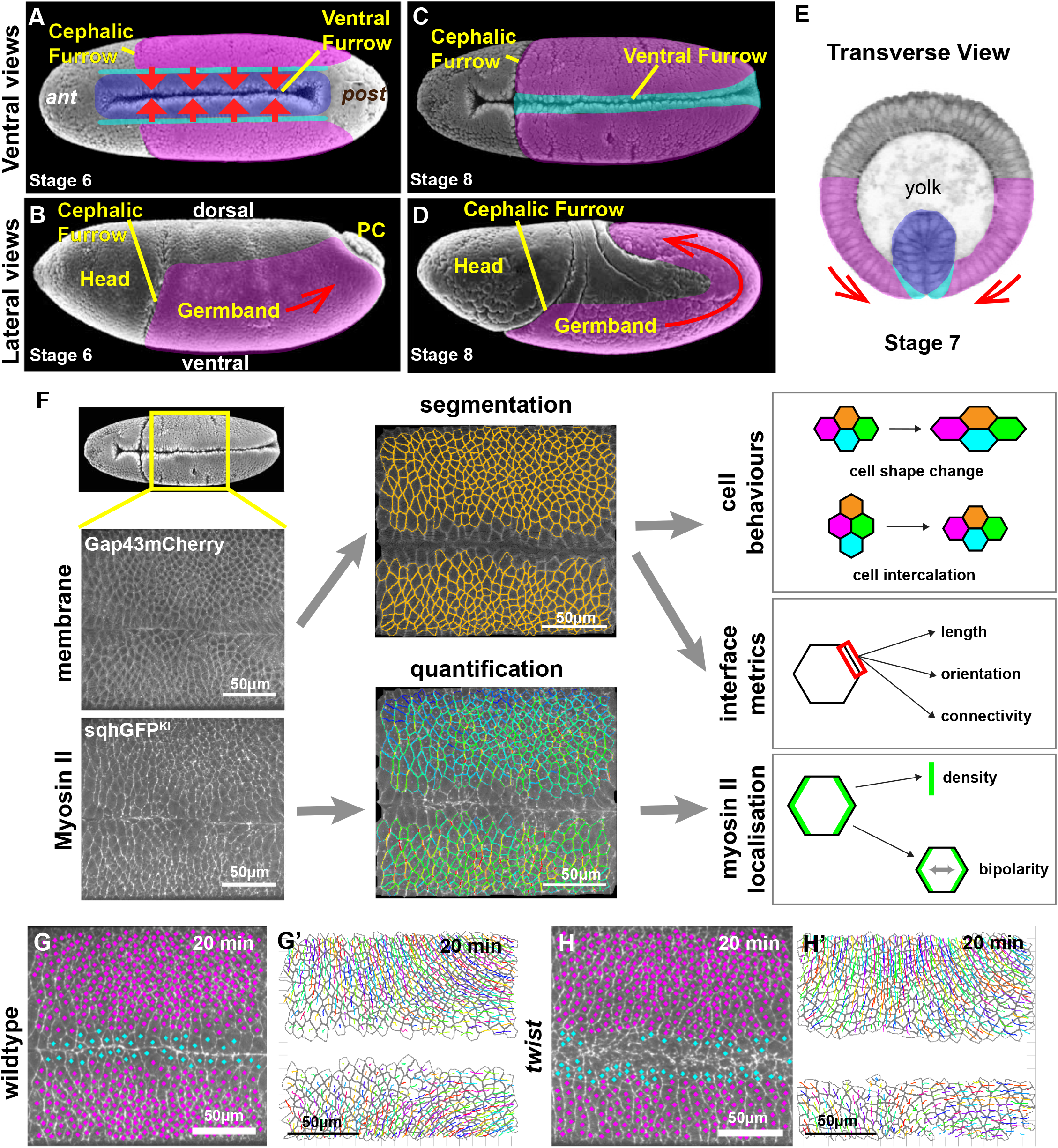
Investigating the impact of mesoderm invagination on germband extension. Scanning Electron (A-D) and tranverse section (E) micrographs from Flybase showing anatomy of *Drosophila* embryos from early to mid germband extension (stages 6 to 8). Coloured overlays highlight germband (magenta), mesectoderm (cyan) and mesoderm (blue). Red arrows depict tissue movements. PC, pole cells; ant, anterior; post, posterior. Anterior to the left in (A-D). (F) Workflow diagram summarising image acquisition and analysis. Panels are maximum intensity projections of live spinning-disc confocal imaging of Gap43Cherry (upper panel) and sqhGFP^KI^ (lower panel), followed by segmentation and tracking of Gap43Cherry channel to extract cell behaviours and interface metrics and quantification of Myosin II channel to extract Myosin density on cell interfaces (blue, low; red, high) and bipolarity measures. (G and H) Example frames for defining cell types in wildtype (G) and *twist* (H) movies (mesoderm/mesectoderm in cyan, ectodermal germband in magenta) overlayed on Myosin II channel (maximum intensity projection). Unmarked cells are those poorly tracked and excluded from the analysis. See also Fig.1-S1.(G’ and H’) Examples of segmentation of ectodermal cells (grey) and tracking of cell centroids (coloured lines) in wildtype (G’) and *twist* (H’), showing trajectories of cells over the previous 5 minutes.

We focus here on the interaction between ectoderm and mesoderm. Mesoderm invagination pulls the germband directly against the direction in which it will undergo convergence, and perpendicular to the direction in which it will extend (Fig. 1A-E). Therefore, from a mechanical perspective, it could act to hinder germband convergence and extension. However, the ectoderm appears to be protected from the mechanical pull of mesoderm invagination due to the behaviour of neighbouring tissues. (Rauzi et al., 2015) report that the neurogenic ectoderm (germband) acts as a ‘stiff non-deformable unit’ and these cells translate en-bloc ventrally in response to mesoderm invagination enabled by the compliance of the dorsal tissue. Additionally, the peripheral cells of the mesoderm have been found to undergo substantial stretch during mesoderm invagination, whilst the ectodermal cells beyond the mesectoderm are described as remaining ‘inert’ (Bhide et al., 2021). This suggests that stretching of the peripheral mesoderm and dorsal tissue accomodates the mechanical tension induced by the invagination of the mesoderm, providing a mechanical buffer for the ectoderm. However, contrary to the possibility that mesoderm invagination might slow down germband extension, or the suggestion that the germband is protected from its mechanical impact, other studies suggest that mesoderm invagination might speed up germband extension (Butler et al., 2009; Farrell et al., 2017; Wang et al., 2020; Streichan et al., 2018; Gustafson et al., 2022). In particular, it has been proposed that the rate of polarised cell intercalation increases in response to mesoderm invagination through a mechanotransduction mechanism leading to enhanced planar polarisation of Myosin II (Streichan et al., 2018; Gustafson et al., 2022).

Some of these discrepancies can be explained by the challenge of quantitatively comparing wildtype embryos with mutant embryos defective for mesoderm invagination (see discussion in Lye et al., 2015). Here we tackle this difficulty by acquiring imaging datasets that are as comparable as possible between wildtype and *twist* mutant embryos. We use these datasets to compare different metrics, including Myosin II densities and junctional behaviours (Fig. 1F). We investigate whether the mesoderm mechanically interacts with the extending ectoderm and systematically evaluate the different hypotheses of how convergence and extension could be impacted by such an interaction. Specifically, a DV-oriented pull by the mesoderm could augment Myosin II enrichment at DV-oriented cell junctions via a mechanotransduction mechanism, thus increasing the rate of junctional shrinkage during neighbour exchange events. Alternatively, if no such mechanotransduction occurred, then as DV-oriented cell junctions stretch in response to mesoderm invagination, cortical Myosin II might become diluted. Additionally, changing the balance of extrinsic forces could have mechanical effects on the speed of junctional shrinkage and growth throughout the process of cell neighbour exchange. Finally, mesoderm invagination might help align cells, and subsequent neighbour exchanges, with the embryonic axes, so that neighbour exchanges drive maximal tissue convergence and extension.

## RESULTS

### Comparing germband extension in wildtype and *twist* mutant embryos

To investigate whether mesoderm invagination has an impact on germband extension (GBE), we carried out a quantitative analysis of germband extension in wildtype and *twist* mutants, which lack mesoderm invagination. We chose to analyse *twist* mutants rather than *snail* or *twist snail* mutants, because although ventral furrow formation fails in all these mutants, some contractility remains in mesodermal cells in *twist* mutants alone, which decreases the width of the mesoderm and makes the ventrolateral field of cells more comparable with wildtype (in *snail* or *twist snail* mutants, no contractility remains and the uninvaginated mesoderm takes more space at the surface of the embryo; Martin et al., 2009).

We acquired movies of the ventral side of embryos from before the onset of mesoderm invagination until the middle of GBE, for both wildtype and *twist* mutants (n= 4 wildtype, n= 6 *twist* embryos throughout this study). We labelled cell membranes with Gap43mCherry, and Myosin II with GFP-tagged knock-in of MRLC to visualise all Myosin II molecules (called *sqhGFP^KI^* here, Proag et al., 2019). Strains were constructed to ensure that these labels were expressed at the same level in wildtype and *twist* embryos (Methods). We also carefully controlled the temperature during imaging. Cell contours were segmented from the *Gap43Cherry* channel and tracked over time (Methods and Movies 1 and 2) to calculate metrics describing key cell and interface behaviours (Fig. 1F)(Blanchard et al., 2009). Myosin II density and planar polarity were quantified from the *sqhGFP^KI^* channel (Fig. 1F)(Tetley et al., 2016).

Our field of view captured the convergence and extension of the ectodermal tissue on the ventrolateral surface of the embryo but also included mesodermal cells and mesectodermal cells. The inclusion of the ventral midline enabled us to precisely measure orientations with respect to this landmark (see Methods). To restrict our analysis to the extending ectoderm, we carefully excluded mesodermal and mesectodermal cells based on: i) whether they had invaginated (wildtype only), ii) their proximity to the midline, and iii) the timing of their cell divisions (Fig. 1G-H’, Fig. 1-S1) (Leptin and Grunewald, 1990; Arora and Nusslein-Volhard, 1992).

Next we checked that the ectodermal cells analysed in wildtype and *twist* are comparable in terms of DV and AP genetic patterning. We examined the expression patterns of key DV patterning genes: *singleminded* (*sim*) is expressed in the mesectoderm, ventral nervous-system defective (*vnd*) is expressed in the ectoderm abutting *sim*, and intermediate neuroblast defective (*i<u>nd</u>*) is expressed next, abutting *vnd* (Reeves and Stathopoulos, 2009). There is some de-repression of both *sim* and *ind* in the uninvaginated mesoderm and the mesectoderm in *twist* mutants, but the expression of all three genes is similar in the ectoderm (Fig. 1-S2). We also checked the expression patterns of the LRR cell surface receptors Tolls 2, 6, 8 and Tartan which are required for the polarised distribution of Myosin II during germband extension downstream of the AP patterning genes (Pare et al., 2019; Pare et al., 2014; Lavalou et al., 2021; Sharrock et al., 2022; Pare and Zallen, 2020). We confirmed that these genes are expressed with the same patterns in *twist* mutants as in wildtype (Fig. 1-S2). Therefore, the analysed regions of the ectodermal germband in wildtype and *twist* mutants can be used to assess the physical effect of mesoderm invagination on germband extension independently of patterning.

To be able to compare averaged metrics between wildtype and *twist* mutants, we synchronised the movies in time, using the start of tissue extension along the antero-posterior axis in each movie as time zero (Methods) (Fig. 1-S3A,B) (Butler et al., 2009; Lye et al., 2015; Sharrock et al., 2022; Tetley et al., 2016). To check that timelines of wildtype and *twist* mutant movies were comparable once synchronised, we checked the timings of two developmental events: when Myosin II becomes detectable apically in the ectoderm and when the first cell divisions occur (Methods). We found that appearance of apical ectodermal Myosin II (around 10 mins before GBE) and the first cell divisions in the lateral ectoderm (around 32 mins after GBE) are grouped in time across movies (Fig. 1-S3C). We note that mesectoderm cell divisions are delayed by nearly 10 mins in *twist* mutants compared to wildtype, which might be a consequence of misregulation of mesodermal and mesectodermal patterning genes. We conclude that our movie synchronisation gives comparable developmental time windows for the two genotypes.

We next determined the number of cells tracked at each time-point in wildtype and *twist* movies (Fig.1-S3D-F). At early time points, we track a total of approximately 800 cells (from 4 movies) for wildtype and a total of 2000 cells (from 6 movies) for *twist*. By the end of the period of analysis, in excess of 2000 cells are included for both wildtype and *twist*.

In summary, the above characterisation demonstrates that we have acquired comparable live imaging datasets of hundreds of tracked cells of the ectodermal germband for wildtype and *twist* embryos covering a developmental window of 15 minutes prior to 30 minutes after the onset of germband extension. We confirmed that DV and AP patterning in the extending ectoderm is unchanged in *twist* mutant compared to wildtype, allowing us to focus on the mechanical impact of mesoderm invagination on germband extension.

### The invaginating mesoderm pulls on the extending ectoderm

As the mesoderm invaginates, cells of the ectodermal germband undergo translation ventrally in wildtype, indicating that these two epithelial tissues are mechanically connected. This movement towards the midline is much reduced in *twist* embryos as expected from the lack of invagination (Fig. 2A, A’). Because of this mechanical linkage, we hypothesised that the invaginating mesoderm must be pulling on the adjacent ectodermal germband, thereby generating a DV-oriented tensile stress. Gradients of cell elongation and apical area increase have been found as a signature of epithelial tissues being subject to tension from extrinsic forces generated by neighbouring morphogenetic movements (Butler et al., 2009; Lye et al., 2015; Bhide et al., 2021; Gustafson et al., 2022). To ask whether the ectoderm is being pulled by the invaginating mesoderm, we analysed these metrics for all ectodermal cells in the field of view for 4 wildtype and 6 *twist* embryos (Methods and Fig. 2B-H).

**Figure 2:**
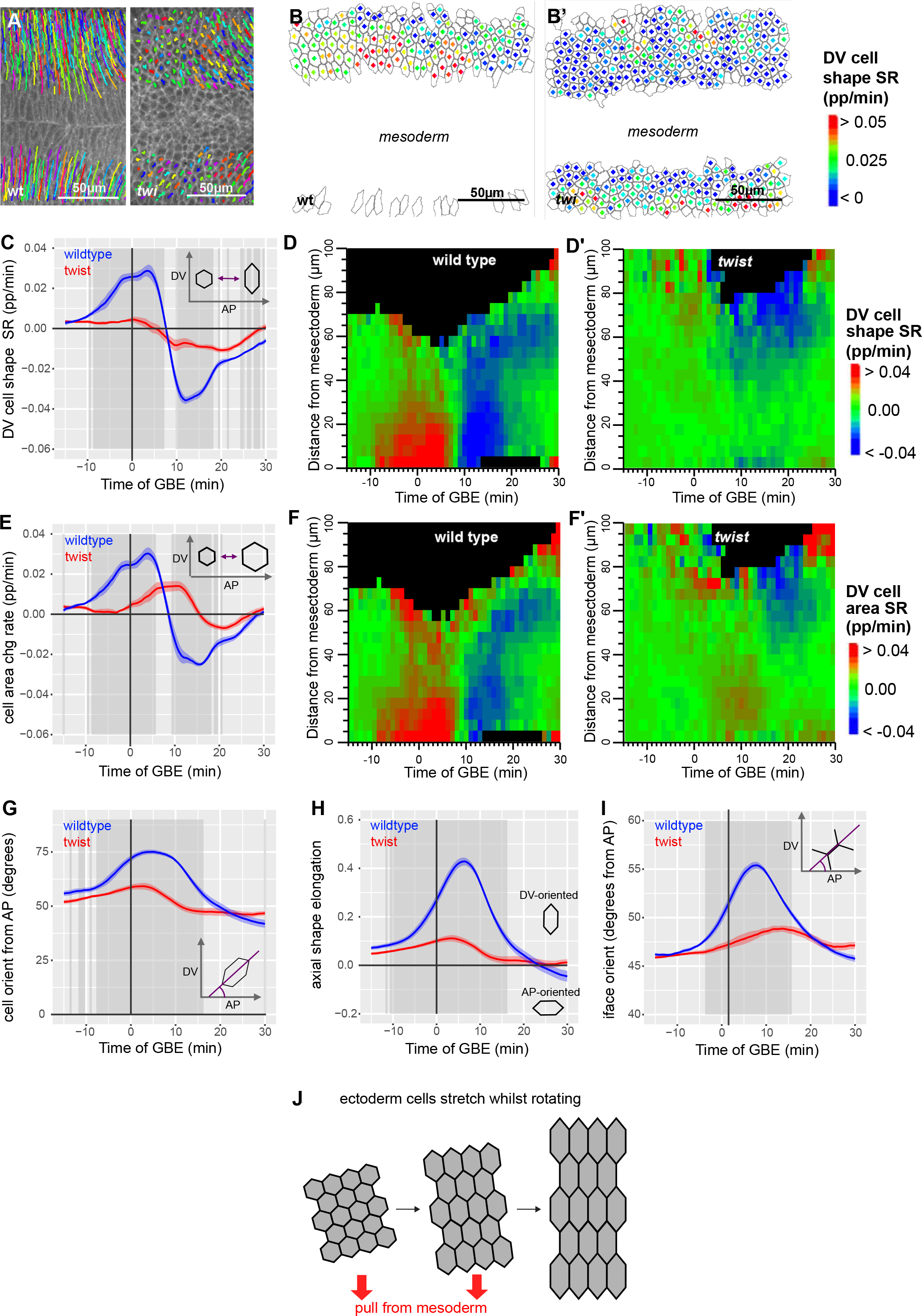
Ectodermal cells during GBE are stretched and rotate in response to ventral pull from mesoderm invagination. (A, A’) Cell trajectories (coloured lines) of germband cells over 5-10 minutes of GBE overlaid on Gap43Cherry image at 10 minutes of GBE (maximum intensity projection). Anterior to the left, ventral centrally. (A) Wildtype (wt): germband cells move significantly ventrally and begin to move posteriorly. (A’) *twist* (t*wi*): germband cells move only slightly ventrally and begin to move posteriorly. (B, B’) Analysis of cell shape change strain rate, projected along DV axis (proportion/minute) in ectodermal cells in wildtype (B) and *twist* (B’) at 2.5 minutes into GBE. Cell shape change of each cell is indicated by coloured square at the centre of each cell (cell outlines, grey). (C) DV-projected cell shape change strain rate (proportion/minute) of ectodermal cells, summarised for wildtype (blue) and *twist* (red) against time of GBE. (D) DV projected cell shape change strain rate (proportion/minute) of germband cells, summarised for wildtype and *twist* against time of GBE and distance from mesectoderm (µm) (see Methods). (E) Cell area change strain rate (proportion/minute) of germband cells, summarised for wildtype (blue) and *twist* (red) against time of GBE. (F) Cell area change strain rate (proportion/minute) of ectodermal cells, summarised for wildtype and *twist* against time of germband extension and distance from mesectoderm (µm) (see Methods). (G) Average orientation of principal axis (degrees from AP axis) of ectodermal cells against time of GBE for wildtype (blue) and *twist* (red). Note that if cells are randomly oriented then the average cell angle would be 45 degrees. (H) Axial shape elongation bias of ectodermal cells against time of GBE for wildtype (blue) and *twist* (red). This measure gives a value of 1 for cells strongly elongated in DV, minus 1 for cells strongly elongated in AP and 0 for isotropic cells or elongated cells at 45 degrees to embryonic axes (see methods). (I) Average orientation of cell interfaces from the AP axis of ectodermal cells for wildtype (blue) and *twist* (red) plotted against time of GBE. Note that if interfaces orientations are not biased to a particular direction then the average orientation should be 45 degrees. Throughout the manuscript dark grey shading indicates periods of significant difference (p<0.01, see Methods) and ribbons show standard error between genotypes. (J) Summary of average ectodermal cell behaviour in response to tensile force from invaginating mesoderm. Cells stretch in DV whilst rotating to become better aligned with the DV embryonic axis.

In wildtype embryos, apical cell lengths along DV (Fig. 2B, C) and apical areas (Fig. 2E) both increase from 15 minutes before the start of GBE, indicating that cells are starting to stretch. The rate of ectodermal cell stretching rapidly increases until approximately 3 minutes into GBE and then rapidly decreases again. Cells continue to stretch in DV until approximately 7.5 minutes into GBE, and then start to shrink in DV (Fig. 2C-E). The period of cell stretching corresponds to when the mesoderm starts contracting and then invaginates in the movies and is consistent with previous reports of cell stretching in the ectoderm in response to mesoderm invagination (Butler et al., 2009; Lye et al., 2015; Gustafson et al., 2022; Farrell et al., 2017). We plotted cell shape changes and apical area changes on spatiotemporal plots to examine how they evolve in time and space. In wildtype, cell shape changes and apical area changes have the same spatiotemporal pattern confirming that the cells are stretching in DV and indicating that the cells are under apical tension (red signal in Fig. 2D, F). Also, there is a gradient in the observed cell stretching from ventral to lateral, suggesting that the source of tension is positioned ventrally, as expected if the mesoderm is pulling. Consistent with this notion, in *twist* embryos, cell stretching in DV is virtually abolished (Fig. 2. C-F). We conclude that the invaginating mesoderm generates a tensile force that is transmitted across the germband causing ectodermal cells to stretch along DV, with a ventral to dorsal gradient.

This tissue level tensile force could also cause ectodermal cells to reorient, so we quantified cell orientation (Fig. 2G, H). First, we plotted cell orientation over time, and find for both wildtype and *twist* that cells are slightly DV-oriented (average angle greater than 45 degrees) already 15 minutes before the start of germband extension (Fig. 2G). This trend increases rapidly in wildtype until about 5 minutes into GBE, reaching a maximum average angle of 75 degrees, showing that cells become very DV-oriented. Cells in *twist* mutants also become briefly more aligned with the DV axis, but to a much lesser extent than wildtype, peaking at about 60 degrees. A combined measure of cell elongation and cell orientation, which we call “axial cell elongation bias” confirms that in wildtype, cells rotate to become more DV-oriented as they stretch in DV (Fig. 2H and Methods). This strong effect occurs from 5 mins prior to 15 mins after the start of GBE (peaking around 7 minutes) and is mostly absent in *twist*, demonstrating that mesoderm invagination not only causes cells to stretch but also to realign perpendicular to the ventral midline. We also plotted average orientation of cell junctions from the AP axis, and find that at -15mins from GBE start, the average orientation is approximately 45 degrees for both wildtype and *twist*, showing no bias in directionality of interfaces at this timepoint (Fig. 2I). In wildtype, this average increases between -10 and 8 minutes as cells stretch in DV, peaking at approximately 55 degrees, indicating a bias towards DV orientation consistent with DV-oriented interfaces becoming more aligned with the DV axis as cells stretch. In contrast for *twist*, the average changes little over time consistent with the finding that there is little DV stretch in cells in *twist* mutants.

Extrinsic tissue-level forces can drive cells to move past each other leading to passive cell intercalations (Caussinus et al., 2008; Blackie et al., 2021). To ask whether the DV-oriented pull of mesoderm invagination causes cell slippage in the ectoderm, we quantified the cell intercalation strain rate as previously (Methods) (Blanchard et al., 2009). We find that the intercalation strain rates along the DV axis is negligible or negative during the period of DV cell stretch, with no difference between wildtype and *twist* mutants (Fig. 2-S1). This demonstrates that the cell stretching caused by mesoderm invagination in the ectoderm is not accompanied by passive cell rearrangement.

In summary, our results show that in wildtype the contraction force from the invaginating mesoderm causes the ectodermal cells of the germband to stretch in DV and change orientation, without causing cells to rearrange along the axis of tension. We next asked whether this mechanical force exerted by the invaginating mesoderm onto the ectoderm impacts on axis extension.

### Comparing Myosin II planar polarisation in wildtype and *twist* mutants

Myosin II-driven polarised cell intercalation is the main cell behaviour causing the convergence and extension of the *Drosophila* germband (Gheisari et al., 2020). During this process of cell neighbour exchange, DV-oriented junctions between AP cell neighbours shrink, so that these cells lose contact with each other, and then a new cell junction grows in the perpendicular orientation (Fig. 3A). Cell junction shrinkage is driven by Myosin II, which is enriched in shrinking junctions under the control of the AP patterning genes. In addition to genetic regulation, there is some evidence in the *Drosophila* germband that Myosin II can be recruited through a mechanosensitive feedback mechanism (Fernandez-Gonzalez et al., 2009; Scarpa et al., 2018). Since mesoderm invagination generates a DV-oriented tensile force within the ectoderm, mechanosensitive recruitment or stabilisation of Myosin II might contribute to the enrichment of Myosin II on DV-oriented cell junctions. Alternatively, cortical Myosin II might be diluted on DV-oriented cell junctions as they are stretched by the invaginating mesoderm. We therefore asked how Myosin II concentration on DV-oriented junctions is affected by mesoderm invagination.

**Figure 3:**
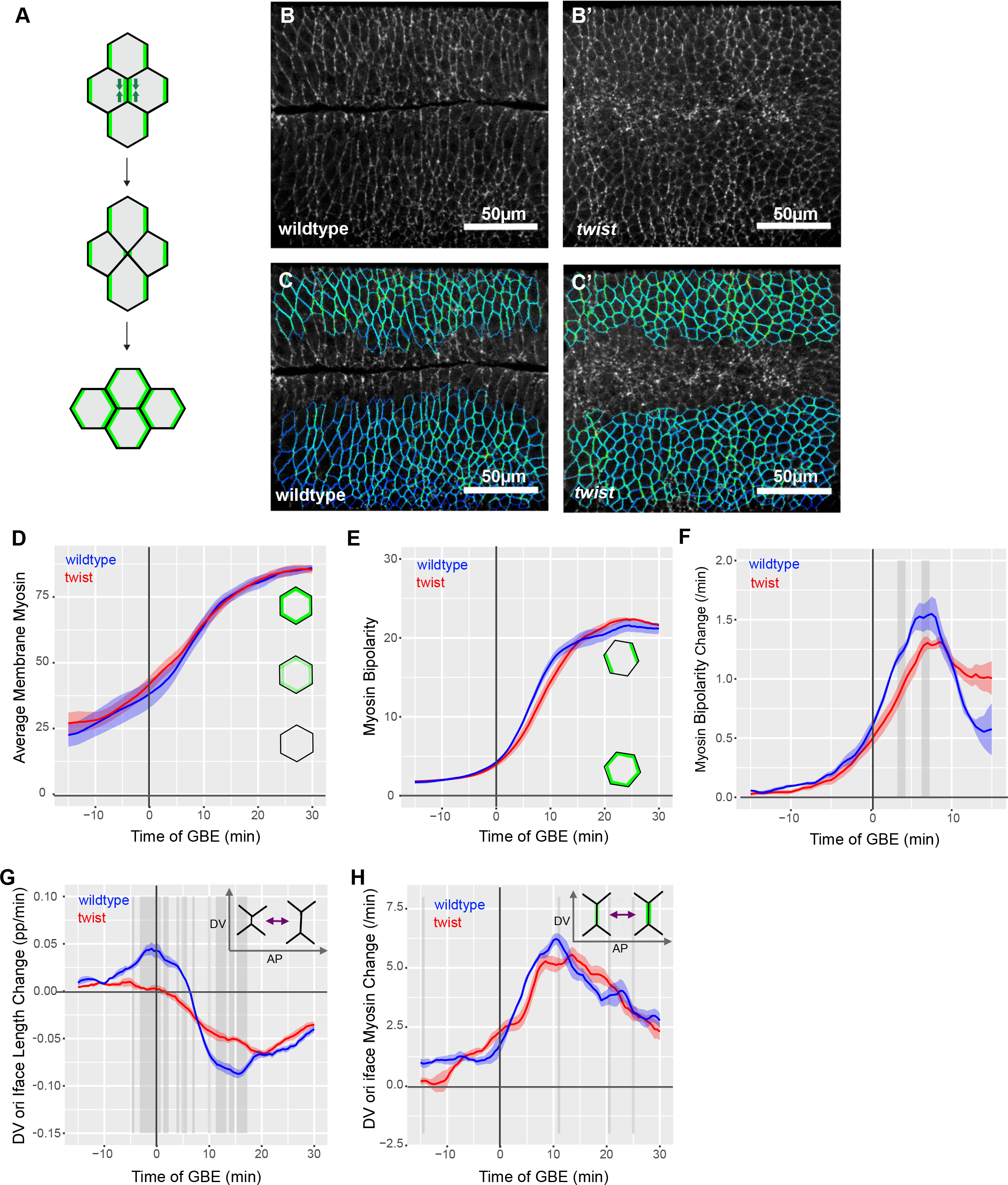
Comparing Myosin II bipolar enrichment in wildtype and *twist* mutants. Diagram showing Myosin II driven cell intercalation. Myosin II (green) shows planar polarised localisation, being enriched in a bipolar manner on the DV-oriented junctions (between cells neighbouring each other along the AP axis). Interface shrinkage (grey arrows), leads to a transient 4-way vertex, followed by growth of a new AP-oriented interface. (B, B’) Normalised Myosin II signal at the level of adherens junctions extracted from confocal image stack for quantification in wildtype and *twist* (B’) at 10 minutes of GBE. See methods for details of extraction and normalisation of signal. (C, C’) Quantified interface Myosin II signal of ectodermal cells corresponding to images in (B and B’) for wildtype (C) and *twist* (C’). Dark blue corresponds to lowest Myosin II density through to red for highest Myosin II density. Note there are no interfaces at this timepoint with highest Myosin II density (red), so green-yellow shows highest Myosin in these frames. See methods for details of quantification. (D) Average density of Myosin II on membranes of ectodermal cells, summarised for wildtype (blue) and *twist* (red) against time of GBE. (E) Unprojected bipolarity of Myosin II of ectoderrmal cells, summarised for wildtype (blue) and *twist* (red) against time of GBE. (F) Rate of change of unprojected bipolarity of Myosin II of germband cells, summarised for wildtype (blue) and *twist* (red) against time of germband extension. (G) Proportional rate of interface length change, for DV-oriented (oriented more than 45 degrees from the AP axis) interfaces only, summarised for wildtype (blue) and *twist* (red) against time of GBE. (H) Rate of Myosin II density change, for DV-oriented (oriented more then 45 degrees from the AP axis) interfaces only, summarised for wildtype (blue) and *twist* (red) against time of germband extension.

First, we quantified fluorescence as a readout of Myosin II density on all cell apical interfaces in the ectoderm, for our wildtype and *twist* movies (Fig. 3B-C’) (Methods). We plotted average fluorescence intensity at all membranes of these cells over time, and observed that in both wildtype and *twist,* fluorescence intensity starts low before germband extension, consistent with our previous observations (Fig. 3D) (Tetley et al., 2016). Membrane Myosin II levels then increase, gradually at first and then more rapidly until approximately 10 minutes after the start of GBE, after which fluorescence intensity continues to increase more slowly. Wildtype and *twist* curves are virtually identical demonstrating that accumulation of Myosin II at apical cell cortices associated with cell membranes is unaffected in *twist* mutants (see also Fig. 1-S3C).

In our movies Myosin II appears enriched on membranes oriented approximately parallel to the DV embryonic axis in both wildtype and *twist*, as expected from the expression pattern of LRR receptors (see above). We quantified this Myosin II planar polarisation by assessing Myosin II bipolar distribution around the membrane of each cell (Tetley et al, 2016). Because there are differences in cell orientation between wildtype and *twist* (see Fig. 2G-I), we plotted the average bipolarity of germband cells, independent of the orientation of the bipolarity. In both wildtype and *twist* embryos, this unprojected Myosin II bipolarity measure starts to increase around the start of germband extension, increases quickly in the first 15 minutes of germband extension and then stays high for the remaining 15 mins, with no significance difference between the two genotypes (Fig. 3E). Spatiotemporal plots for these data show that these temporal patterns are similar for all cells in the field of view and that there is no obvious ventral to dorsal gradient in wildtype (Fig. 3-S1). Although there is no significant difference in unprojected bipolarities at any time between the two genotypes, the *rate* of increase appears faster between 0 and 10 minutes of GBE in the wildtype compared to *twist* mutants (Fig. 3E). Therefore we calculated the rate of change of Myosin II bipolarity, and we found that in *twist* embryos it is slightly reduced compared to wildtype, showing some statistically significant differences around 5 minutes of GBE (Fig. 3F).

Because these differences are seen while cells are stretching (see Fig. 2C-F), we asked whether cell interface stretching is associated with an increased rate of Myosin II recruitment. We plotted the proportional rate of stretching of DV-oriented interfaces and saw that significant stretching occurs in wildtype, but not in *twist,* between approximately -10 and 7 minutes of GBE, as expected as wildtype cells are stretching in DV over this period of time (compare Fig. 3G with Fig.2C). We then plotted the rate of Myosin II density change for DV-oriented interfaces, and observed that the rate of Myosin II recruitment increases rapidly during this time period in wildtype, but also in *twist* mutants whose DV-oriented interfaces undergo very little stretching (Fig. 3H). Shortly after the period of interface stretching, at approximately 11 minutes in GBE, there is a briefly significant period of time when the rate of recruitment is higher in wildtype than in *twist*. Altogether, this indicates that if there is any contribution of mechanotransduction to Myosin II localisation early in GBE in response to the physical pull by the mesoderm on the ectoderm, it is at the limit of what we can detect with our measures. Our results also show that Myosin II density is maintained in DV-oriented junctions that are stretching under tension from mesoderm invagination, compared to unstretched DV-oriented junctions in *twist* embryos, indicating that a homeostatic mechanism maintains Myosin II cortical density despite significant anisotropic cell stretching.

### Comparing junctional shortening speed and rate of neighbour exchange in wildtype and *twist* mutants

In addition to intrinsic forces within the germband driving its convergence and extension, ectoderm extension is facilitated by an extrinsic pull (along AP) from the posterior endoderm (Butler et al., 2009; Collinet et al., 2015; Kong et al., 2017; Lye et al., 2015) Both the convergence and the extension in the germband could be mechanically counteracted by the orthogonal pull (along DV) from the invaginating mesoderm. If so, in the absence of mesoderm invagination (*twist* mutants), we might expect junctional shortening to speed up due to the lack of a DV tension opposing junctional shrinkage. Thus, boundary conditions imposed by the endoderm and mesoderm invaginations could affect the behaviour of ectodermal cells during germband extension. Junctional shortening rates could also be influenced by mesoderm invagination in the opposite way through mechanosensitive Myosin II recruitment, as considered above.

We examined whether mesoderm invagination impacts on the rate of junctional shrinkage of the extending ectoderm throughout early and mid germband extension (0-30 mins). We find that the proportional rate of junctional shrinkage is not significantly different between wildtype and *twist,* but there is a trend for proportional contraction rates to be higher in wildtype interfaces (Fig. 4A). Assuming a viscous constitutive law for the junctions over minute timescales, proportional shrinkage rate should be directly proportional to mechanical stress. We therefore quantified the density of Myosin II specifically on shrinking interfaces (Fig. 4B), since Myosin II is proposed to be the main source of mechanical stress driving junction shrinkage (Bertet et al., 2004; Zallen and Wieschaus, 2004). We find that wildtype Myosin II densities tend to be higher than those of *twist* mutants, but with only a very brief period of statistically significant difference between the two genotypes, at 5 minutes prior to neighbour exchange. This is consistent with the very mild effect on Myosin II in response to mesoderm invagination described in the section above. The slight decrease in Myosin density on shrinking interfaces in *twist* mutant could explain the slight decrease in their proportional contraction rate.

**Figure 4:**
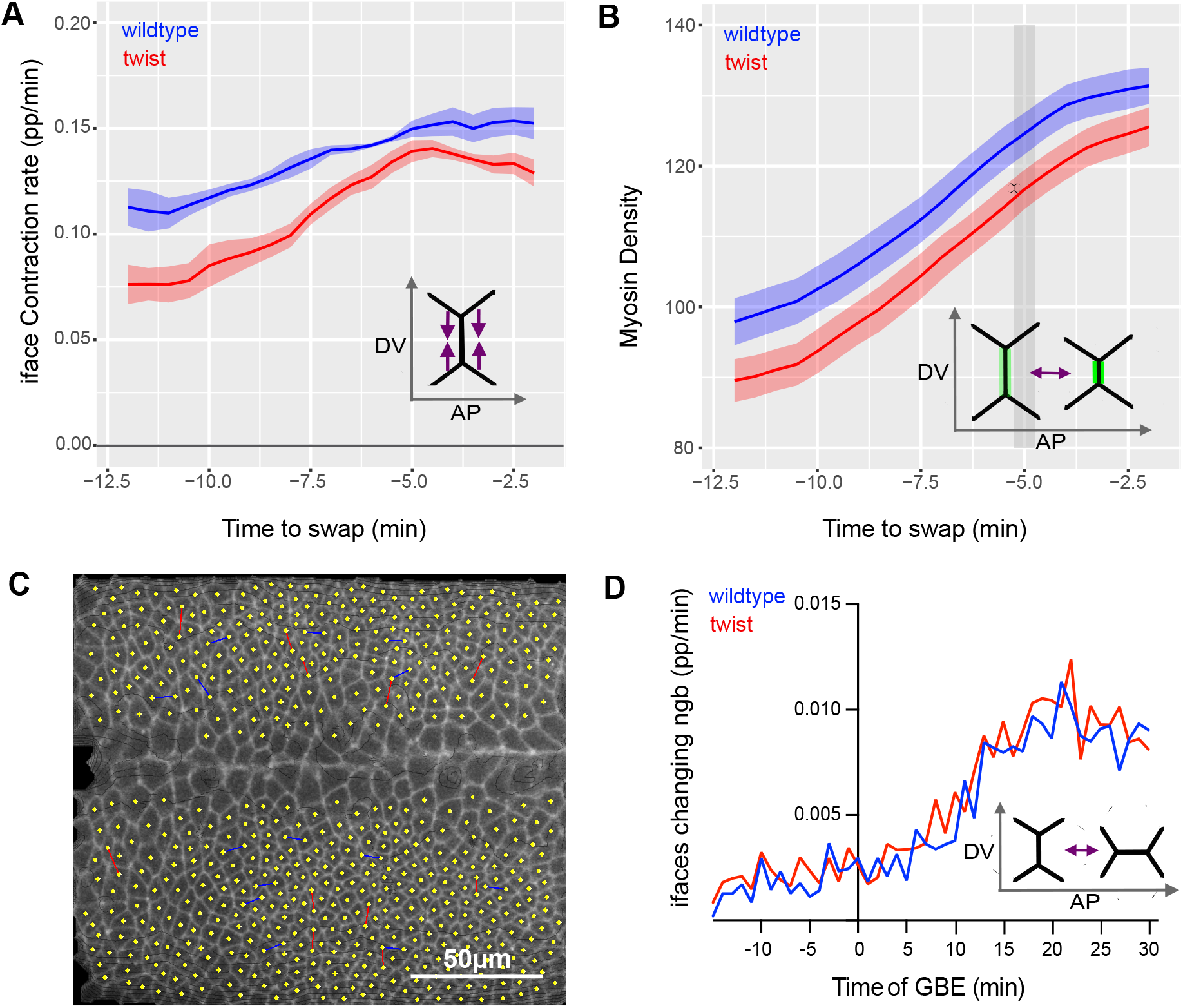
Comparing junctional shortening speed and rate of neighbour exchange in wildtype and *twist* mutants. (A) Proportional rate of interface shrinkage plotted against time to swap during GBE (data for 0-30 minutes of germband extension) summarised for wildtype (blue) and *twist* (red). (B) Average density of Myosin II on shrinking interfaces, plotted against time to an intercalation event (swap) during GBE (data for 0-30 minutes of germband extension) summarised for wildtype (blue) and *twist* (red). (C) Example of detection of neighbour exchange events in the germband in wildtype at 20 minutes of GBE. Loss of neighbours is shown as blue lines between cell centroids, whilst gain of neighbours is shown in red. Cell centroids are shown in yellow. Underlaid is an image of Gap43Cherry signal at level of adherens junctions extracted for tracking. (D) Rate of neighbour exchange. Number of interfaces involved in a T1 swap expressed as a proportion of the total number of DV-oriented interfaces for all tracked ectodermal germband cells, summarised for wildtype (blue) and *twist* (red) over time of germband extension.

The speed in junctional shortening should influence the rate of polarised cell intercalation, so we also looked at a measure of the latter. We tracked individual neighbour exchange events (swaps in junctional connections between cells, see Methods) over time (Fig. 4C). We find that both wildtype and *twist* embryos show a comparable proportion of interfaces exchanging neighbours throughout the course of germband extension with no discernible delay in *twist* compared to wildtype (Fig. 4D). We also find that the majority of neighbour exchange events occur after 15 minutes of germband extension.

In summary, we conclude that there is no effect of mesoderm invagination on the speed of junction shrinkage. Consistent with these observations, we find no difference in the proportion of cells undergoing intercalation events between wildtype and *twist* mutants.

### Comparing junctional elongation speed and AP cell elongation in wildtype and *twist* mutants

Following shrinkage of Myosin II-enriched cell junctions, cells exchange neighbours and cells previously separated from each other along the DV axis come into contact, forming a new junction (approximately AP-oriented) to drive extension of the tissue along the AP axis (Collinet et al., 2015). As for junctional shortening speed, the balance of boundary conditions on the extending ectoderm (posterior pull by endoderm invagination and counteracting orthogonal pull by mesoderm invagination) could impact junctional elongation speed. Specifically, an increase in junctional elongation rates might be expected as new AP-oriented junctions would grow in the absence of an orthogonal tension (along DV) that could slow junction growth. To address this, we i) compared new junction growth in *twist* mutant and wildtype embryos and ii) examined AP cell elongation rates, which would be both predicted to be affected by the change in boundary conditions.

First, we quantified the speed of growth of new junctions in wildtype and *twist* and found that new junctions grow quicker in *twist* than in wildtype (Fig. 5A). We propose that in wildtype the DV tension in the germband that is induced by mesoderm invagination slows the rate of elongation of new junctions. In addition to cell intercalation, AP cell elongation (AP cell shape strain rate) has been shown to contribute to germband extension and this is driven by AP tension induced by the invagination of the posterior endoderm (Butler et al., 2009; Lye et al., 2015). Cell stretching in either AP or DV requires apical area increase, and this may have an energetic cost due to the bulk properties of the cell. We indeed can verify that cell area change is globally higher in wildtype than in *twist* due to the ventral pull from mesoderm invagination (Fig. 5-S1B,B’). Therefore, the removal of the DV-oriented force in *twist* mutants may lead to an increase in AP cell elongation as cells are not also stretched in the DV direction. Because AP cell elongation rates vary along the AP axis, as they are influenced by both endoderm invagination at the posterior and cephalic furrow formation at the anterior (Butler et al., 2009; Lye et al., 2015), we plotted AP cell elongation rates against time of germband extension and AP position (Fig. 5C, C’). In wildtype, we can see cell shape changes associated with cephalic furrow formation and posterior midgut invagination, with only low levels of AP cell elongation in between. In *twist,* the cell shape changes associated with cephalic furrow formation are less clear, but those associated with posterior midgut invagination are very clear. This difference compared to wildtype is probably due to slight differences in precise AP positioning of embryos within the field of view. More interestingly, the rate of AP cell elongation in the central region of the embryo appears higher in *twist* mutants than in wildtype. To ask whether this difference was significant we plotted AP cell elongation rate against time in wildtype versus *twist* movies, with only this central region included in the analysis (see Methods), and we confirm that AP cell elongation is higher in this region in *twist* mutants than in wildtype, with several bursts of significance (Fig. 5B). A similar analysis for the whole of the imaged region is shown in Fig. 5-S1A.

**Figure 5:**
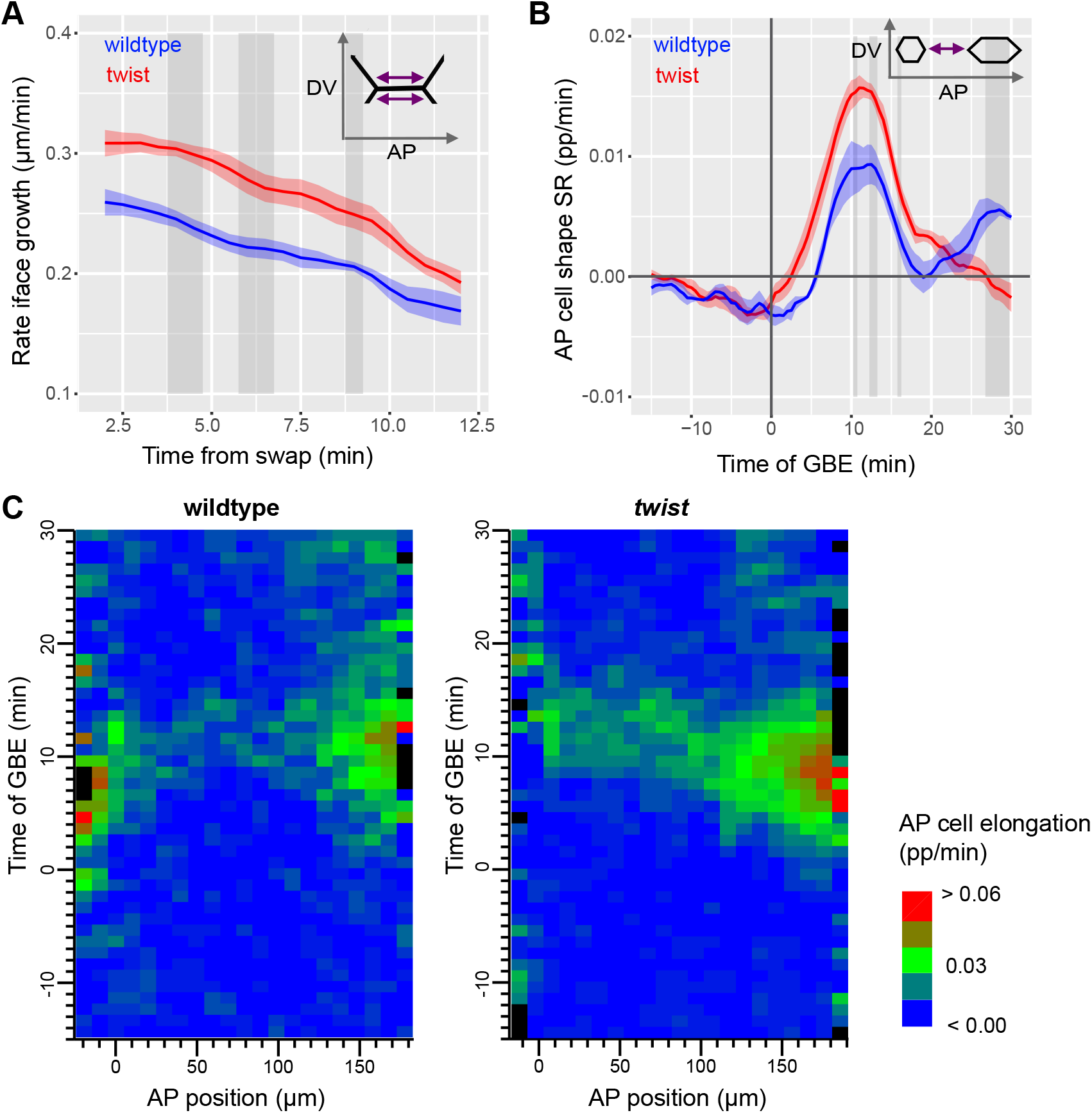
Comparing junctional elongation speed and AP cell elongation in wildtype and *twist* mutants. (A) Rate of interface growth of interfaces in ectodermal germband cells for wildtype and *twist* over time from neighbour exchange events. (B) AP cell shape change strain rate plotted time of GBE summarising data from the central region (avoiding areas of high cell shape change associated with cephalic furrow formation and posterior endoderm invagination) of wildtype (blue) and *twist* (red) movies. (C, C’) Spatiotemporal plots of AP cell elongation plotted against AP position and time of GBE summarising wildtype (C) and *twist* (C’) movies.

In summary, we conclude that a change in the DV boundary conditions in *twist* mutants leads to an increase in the speed of both elongation of new junctions following T1 transitions, and AP cell elongation due to a reduction in DV tension.

### Comparing the orientation of cell interfaces during cell intercalation in wildtype and *twist* mutants

Finally, we asked whether the observed changes in the alignment of cells and cell junctions with the embryonic axes in *twist* compared to wildtype could impact tissue extension, since this could lead to polarised cell intercalation being less well aligned with the embryonic axes, which might lessen the effectiveness of convergence and extension.

First, we examined Myosin II bipolarity alignment with the embryonic axes in *twist* mutants compared to wildtype. We projected our bipolarity cell measure along the anterior-posterior axis to assess the extent to which Myosin II is enriched at the anterior and posterior of each cell (Fig. 6A, Fig. 6-S1). We find that the onset of AP-projected Myosin II polarity is delayed by approximately 5 minutes and significantly reduced in *twist* compared to wildtype, until approximately 10 minutes into GBE. This difference is much more pronounced than for the unprojected data (compare Fig. 3E with Fig. 6A and Fig. 3-S1 with Fig. 6-S1), showing that the Myosin II bipolarity of cells is less well aligned with the embryonic axes early in germband extension in embryos in which mesoderm invagination is defective. We conclude that as well as having very slightly lower rates of Myosin II recruitment (see earlier), the Myosin II enriched junctions in *twist* mutants are now less well aligned perpendicular to the AP axis than in wildtype.

**Figure 6:**
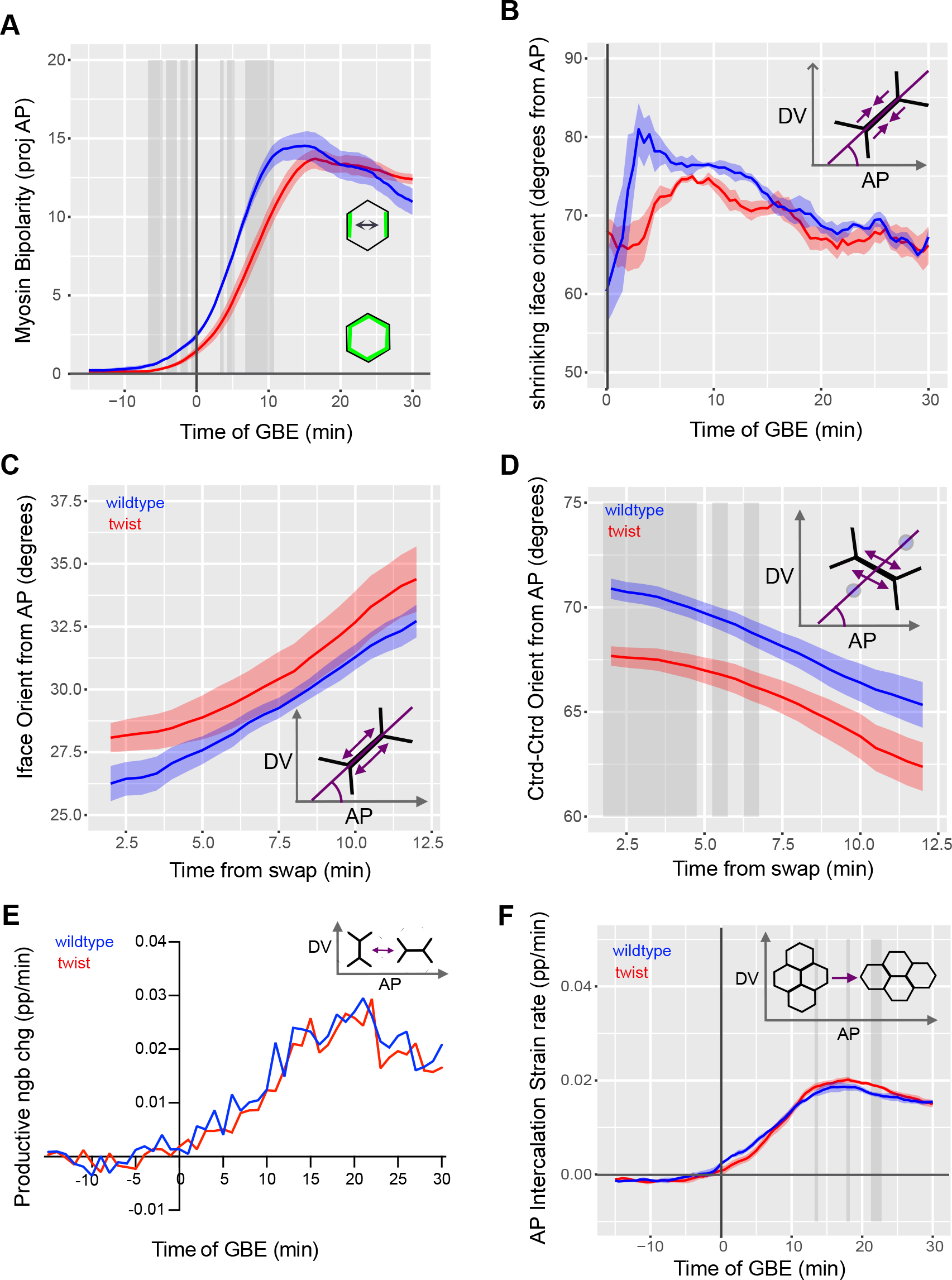
Comparing the orientations of growing and shrinking junctions in wildtype and *twist* mutants. (A) Myosin bipolarity projected along AP of all tracked ectodermal cells, summarised for wildtype (blue) and *twist* (red) plotted against time of GBE. (B) Orientation (degrees from AP-axis) of shrinking interfaces at 6.5 minutes prior to swap of all tracked ectodermal germband cells, summarised for wildtype and *twist* over time of GBE. (C) Orientation (degrees from AP-axis) of growing interfaces in ectodermal germband cells, summarised over the first 30 minutes of germband extension for wildtype and *twist* plotted against time from neighbour exchange event. (D) Orientation (degrees from AP axis) between cell centroids of pairs of ‘new neighbours’ in ectodermal germband cells, summarised over the first 30 minutes of GBE for wildtype and *twist* plotted against time from neighbour exchange event. (E) Rate of productive neighbour exchange. Rate of net gains along the AP axis, using a continuous angular measure for productivity of T1 swaps to axis extension (see Methods), expressed as a proportion of the total number of DV-oriented interfaces for all tracked ectodermal cells, summarised for wildtype (blue) and *twist* (red) over time of GBE. (F) AP-projected intercalation change strain rate (proportion/minute) of all tracked ectodermal cells, summarised for wildtype and *twist* plotted against time of germband extension.

Whilst the number of intercalation events is not reduced in *twist* compared to wildtype (Fig. 4F), the changes in average cell junction orientation and AP-projected bipolarity of Myosin II suggests that shrinking interfaces involved in T1 transitions may be poorly aligned with the DV axis. Therefore we measured the orientation of shrinking interfaces shortly before neighbour exchange throughout germband extension (Fig. 6B). We find no statistically significant difference between the orientations of shrinking interfaces in wildtype and *twist*, but early in germband extension there is a trend for shrinking interfaces to be better aligned with the DV-axis in wildtype, with the difference in orientations not exceeding 15 degrees. The timing of the observed difference in orientations of shrinking interfaces early in germband extension (before 20 minutes GBE) reflects the timing of differences in cell orientations and orientation of the interface population as a whole, suggesting that differences in orientations of shrinking interfaces are due to differences in cell orientations in the germband at this time (see Fig. 2). The fact that we only detect very minor misalignment of Myosin II enriched shrinking interfaces early in GBE suggests that the orientation of shrinking interfaces is not predominantly controlled by the global tissue movements and tension induced by mesoderm invagination.

Because shrinkage of these Myosin II-enriched, DV-oriented interfaces will directly contribute to convergence of the tissue in the DV, we asked whether this minor misalignment of shrinking interfaces had an impact on DV intercalation strain rate. Because the shrinking junctions are to be misaligned with the DV axis by up to 15 degrees early in germband extension, then the *maximum* expected difference in strain rate projected along the DV axis from this misalignment can therefore be estimated as 1 – cos(15) = 3.4%, predicting a very minor effect. Consistent with this, we found no significant reduction in DV intercalation convergence rate in *twist* embryos compared to wildtype embryos (Fig. 2-S1). Previously, in other mutants affecting the orientations of shrinking interfaces, we had seen that the angle between junction growth and shrinkage was not affected, so growing junctions were also misoriented (Finegan et al., 2019). Therefore, since we see some differences in the orientations of shrinking interfaces, we also assessed the orientation of new junction growth. We did not detect a statistically significant difference in orientation of growing interfaces in *twist* mutants compared to wildtype, although growing interfaces do tend to be slightly less well aligned with the AP axis (Fig. 6C). Also, when plotting the angles between the centroids of newly neighbouring cells, we do see that these centroid-centroid angles are significantly less well aligned with the DV axis in *twist* mutants than in wildtype for approximately 6 minutes after a neighbour exchange (Fig. 6D). Therefore, although T1 neighbour exchanges occur at normal rates in *twist* mutants (Fig. 4D), the resulting cell packing is very mildly perturbed compared to wildtype soon after the neighbour exchange, but this effect is shortlived. We also quantified the rate of productive neighbour exchanges contributing to tissue extension (that is net T1 gains along the anterio-posterior axis, taking into account the varying angles of T1 events with respect to the embryonic axes, see Methods) and observe that it is very similar in *twist* compared to wildtype (Fig. 6E). This shows that minor observed differences in angles of T1 events between wildtype and *twist* do not impact on the ability of T1 events to extend the ectodermal germband.

Finally, we measured the intercalation strain rate in AP to ask whether the minor differences that we observe in junction growth speed and orientation cause a difference in the extension of the tissue through intercalation (Fig. 6F). We find no convincing difference between wildtype and *twist* embryos though there is a trend for *twist* intercalation strain rate to be slightly higher, with several short bursts of significant differences between wildtype and *twist*.

In summary, although there are measurable differences in orientation of shrinking and growing interfaces, the effect on tissue extension is negligible. Considering all our results together we conclude that the cell intercalations required for germband extension are not significantly augmented by mesoderm invagination, but they are robust to the extrinsic force that it exerts.

## DISCUSSION

We set out to investigate whether mesoderm invagination impacts on germband extension in *Drosophila* embryos. We find that despite mesoderm invagination pulling and transiently deforming the adjacent ectoderm, the extension of ectoderm is remarkably unaffected by this mechanical interaction. This reveals the robustness of germband extension to a perpendicular tensile stress.

### Mechanical impact of mesoderm on the ectoderm

Previous studies looking at the mechanical impact of mesoderm invagination have focused on cell deformation within the mesoderm domain (Bhide et al., 2021; Fuentes and He, 2022). These studies have shown that while the central region of the mesoderm undergoes apical constriction, a region of about three cells on either side (the most lateral of those being the mesectoderm cells) do not constrict but instead stretch along DV. Beyond the mesectoderm cells, the lateral ectoderm cells were thought to be “inert” and mechanically buffered by the stretching of compliant tissue at its ventral (and also dorsal) edge (Bhide et al., 2021; Fuentes and He, 2022; Rauzi et al., 2015). In this paper, we show that ectodermal cells do respond mechanically to mesoderm invagination. We detect significant cell stretching in the lateral ectoderm, which is lost in *twist* mutants. This suggests that forces generated by the contractile activity of the apical actomyosin in the mesoderm do transmit to the ectoderm via the apical adherens junctions. When mesoderm invagination begins, the level of apical Myosin II is still very low in the lateral ectodermal cells (Fig. 3D), which might explain why ectodermal cells stretch significantly under tension from the invaginating mesoderm, relieving the mechanical stress. We also show that ectodermal cell stretching is not accompanied by cell slippage.

### Mechanisms regulating Myosin II density at the cortex and consequences for cell intercalation

The stretching of ectodermal cell-cell junctions could trigger a mechanosensitive response leading to an increase in Myosin II enrichment, or alternatively, a dilution of Myosin II (Fig. 7A). We find a clear absence of dilution, suggesting that a homeostatic mechanism is able to maintain Myosin II density at the cortex upon rapid stretching. Alternatively, this response might simply be that the AP patterning system controlling Myosin II enrichment on DV-oriented interfaces controls the density of Myosin II at the cortex, rather than the total amount. The result of maintaining Myosin II density in stretching DV-oriented junctions is that they still undergo shrinkage to initiate neighbour exchange events at the appropriate rate, and so mesoderm invagination does not negatively impact this first step of cell intercalation. In contrast, in the case of the peripheral mesoderm, the low levels of apical medial Myosin II seem to be diluted and easily overcome by the pulling forces of the central mesoderm leading to extreme apical stretching of cells, rather than homeostatic maintenance of the actomyosin (Bhide et al., 2021). This highlights the fact that responses of cells to apical tension are stage- and cell type-specific.

**Figure 7:**
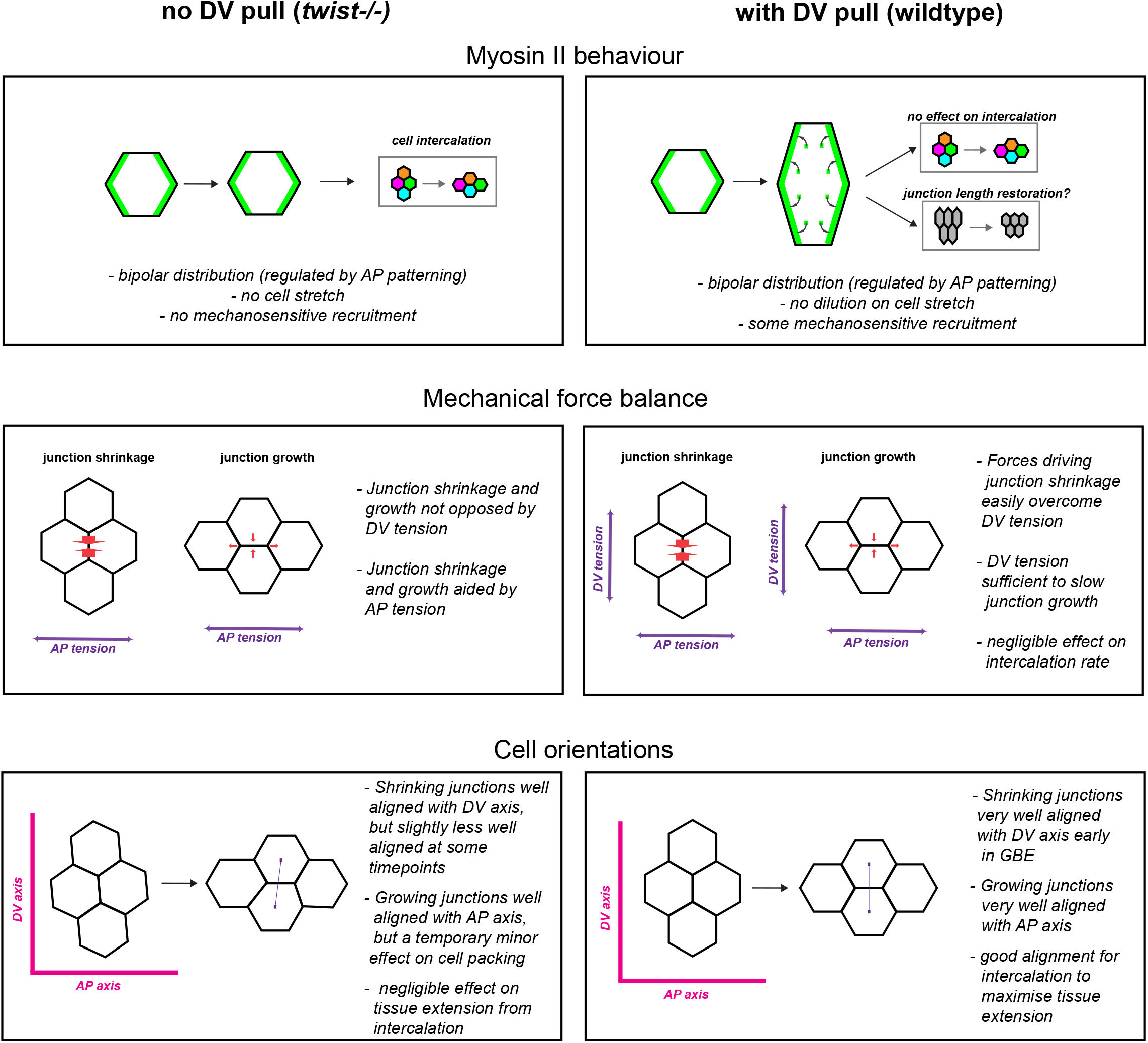
Summary of the consequence of mesodermal pull on cell and cell junction behaviours associated with cell intercalation during germband extension. Left column considers the behaviours in the absence of the mesodermal pull (i.e. *twist* mutant embryos) and right column considers the behaviours in the presence of the mesodermal pull (i.e. wildtype embryos).

We found evidence of only a very mild Myosin II mechanotransductive response; Myosin II recruitment rate was temporarily marginally higher in wildtype than *twist* but this did not have an effect on the shrinkage of cell junctions during neighbour exchange. Our results are in contrast to some other studies which have reported a stronger effect of mesoderm invagination on Myosin II localisation and germband extension, but there are inconsistencies in results between studies, perhaps due to the challenge of acquiring comparable datasets between wildtype and mutants, so the precise effect of mesoderm invagination on germband extension has not been clear (Butler et al., 2009; Farrell et al., 2017; Wang et al., 2020; Streichan et al., 2018; Gustafson et al., 2022; Lye et al., 2015). Mechanosensitive feedback in Myosin II recruitment has been reported at cable-like actomyosin enrichments spanning several junctions connected to each other later in germband extension and at parasegmental boundaries due to linked junctions pulling on each other (Fernandez-Gonzalez et al., 2009; Scarpa et al., 2018). In both cases, a modest decrease in Myosin II recruitment was measured (less than 10%) when the “cables” were cut by laser dissection to decrease tension in the linked junctions. Considering that mesoderm invagination pulls on the ectoderm when apical actomyosin is very low and just starting to increase (Fig. 3D), it is perhaps unsurprising that any mechanosensitive increase in Myosin II would be challenging to detect.

### Restoration of DV cell length after being stretched by mesoderm invagination

Following mesoderm-induced DV cell and cell junction stretching, cells contract to regain an approximately isotropic shape. When comparing rates of junctional shrinkage leading to neighbour exchange, we plotted the proportional rate of length change. Therefore, because wildtype DV oriented cell junctions have been stretched, in absolute terms wildtype shrinking junctions shrink faster than those in *twist* mutants, as cell junctions are both restoring their length after stretching and undergoing shrinkage for intercalation events simultaneously. The transient marginal increase in the rate of Myosin II recruitment in wildtype compared to *twist* (Fig. 3) may be due to a mechanotransductive mechanism that acts to restores junctional length after stretching. Other mechanisms might also contribute to the restoration of cell shape after stretching including their material properties, the activity of medial myosin networks at the apices of cells and three-dimensional constraints to their shape. Nevertheless, the possibility remains that mechanosensitive recruitment or stabilisation of Myosin II on the cortex of stretched junctions is important for cell junctions to resist overstretching and subsequently restore their length, but not for helping to drive cell intercalation as we show here.

### Boundary conditions during germband extension and impact on junction shortening and elongating rates

Loss of mesoderm invagination would be expected to change the boundary conditions during germband extension. The extending germband is pulled along AP as a consequence of endoderm invagination and mesoderm invagination provides a pull along DV. These two pulls can be considered as boundary conditions acting in perpendicular directions on the extending germband. Even after mesoderm invagination is complete, the DV tension may remain as the mesectodermal cells meet and adhere to each other at the midline which could hold tension in the ectoderm. This DV-oriented force could counteract both the AP pulling force and forces generated within the germband promoting convergence and extension such as those generated by Myosin II contraction at shrinking interfaces. If so, the rate of junction shortening and elongation would increase in the absence of mesoderm invagination. We find a clear increase in the rate of junction elongation, and also cell elongation along AP, in *twist* mutants. This suggests that mesoderm invagination does act as a counteracting force. However, despite the increase in junctional elongation rate seen in *twist* mutants, we see little impact of this on the overall rate of cell intercalation.

We do not, however, detect a similar increase in the rate of junctional shortening in *twist* mutants. Based on the above, it is plausible that an increase might be cancelled out by a decrease in junctional shortening linked to a decrease in Myosin II mechanosensitive recruitment in absence of cell stretching. In this model, the opposing force of mesoderm invagination to ectoderm extension would be balanced by the increased contractility of the stretched interfaces. Alternatively, the DV tension generated by mesoderm invagination might just have negligible mechanical effect on the speed of junctional shortening, if the force driving junctional shortening is much greater than that of the DV tension.

### Junction orientation and tissue extension

Finally, mesoderm invagination might impact ectoderm extension by helping to align the junctions of rearranging cells along the main embryonic axes. We do find a significant effect of mesoderm invagination on cell and junction orientation up to approximately 15 minutes of GBE, but relatively few neighbour exchanges occur before this time. Furthermore, the effect on the shortening cell junctions themselves is marginal, even within this first 15 minutes of GBE, suggesting that their orientation is relatively insensitive to tissue-level forces. This could be explained by the rapidly increasing Myosin II planar polarisation of the ectoderm during this period. We and others have shown that Myosin II-enriched interfaces form supracellular cable-like structures that straighten throughout the course of GBE (Fernandez-Gonzalez et al., 2009; Tetley et al., 2016; Sharrock et al., 2022). Thus, this tissue-autonomous straightening might explain why the orientation of shortening junctions is mostly unaffected by mesoderm invagination.

We also found only marginal differences in the orientation of growing junctions between wildtype and *twist* mutants. This is expected since junctions grow in an orientation perpendicular to shortening junctions and this angle is tightly constrained (Finegan et al., 2019). Also, invagination of the posterior endoderm (which still occurs in *twist* mutants) will act to pull growing junctions to be parallel with the AP axis (Collinet et al., 2015). Overall, we find that the minor differences we observe in the orientation of shrinking and growing interfaces do not have an impact on the productivity of cell intercalation events for tissue extension.

In summary we find that despite minor differences in Myosin II recruitment speed, speed of cell junction growth and cell junction orientation between wildtype and twist embryos, the invagination of the mesoderm does not significantly hinder cell behaviours underlying germband extension, nor does it significantly enhance them through mechanosensitive localisation of Myosin II. Therefore, the morphogenesis of germband is robust despite a mechanical pull orthogonal to the direction in which it needs to extend.

## METHODS

### Fly Maintenance and Strains

We used a CRISPR knock-in line (sqh-eGFP[29B], referred to in this work as sqhGFP^KI^) to tag endogenous Myosin II Regulatory Light Chain with GFP; specifically (Proag et al., 2019). We used Gap43mCherry (on the X) to mark cell membranes (Izquierdo et al., 2018). Recombinant chromosomes were made using standard *Drosophila* genetics. *twist* homozygous embryos were collected from sqhGFP29B.Gap43Cherry;twi1/Cyo stocks and wildtype from sqhGFP29B.Gap43Cherry; Gla/Cyo stocks. Standard fly husbandry was used to maintain stocks.

### HCR and antibody staining

HCR and immunostaining methods are as described previously (Sharrock et al., 2022). Stage 5 to 8 embryos were collected from twi1/CyO stocks. Morphology of the ventral furrow and ectopic expression of ind or sim within the mesoderm was used to identify mutants. The HCR probe sets were designed by Molecular Instruments. Primary antibodies used were mouse anti-phospho-Tyrosine (pTyr) (Cell signaling #941) (1:1000) and rabbit anti-phospho-Histone3 (pHis) (Cell signalling #9701; 1:200). Secondary antibodies conjugated to fluorescent dyes were obtained from Jackson ImmunoResearch Laboratories, Invitrogen and Life Technologies. Streptavidin with Alexa Fluor 405 conjugate was from ThermoFisher Scientific.

Embryos were imaged on an inverted SP8 Confocal Microscope (Leica Systems), with a 40X 1.3NA oil-immersion objective. Either a PMT or HyD detector was used alongside a 405/488/546/647nm laser line. Image stacks of 1um Z separations were captured using the Leica Application Suite X Software.

### Movie acquisition

Embryos were mounted ventral/ventrolaterally and a central region of the embryo was imaged on a spinning disc confocal microscope as previously (Tetley et al., 2016). We chose this field of view to avoid the cephalic furrow and the posterior of the embryo, where cells are known to stretch in AP in response the cephalic furrowing and posterior midgut invagination respectively (Butler et al., 2009; Lye et al., 2015). Z-stacks for each channel were collected sequentially. Embryos were imaged every 30 seconds from approximately mid-cellularisation (stage 5) until the onset of cell divisions within the field of view (approximately stage 8) at a controlled temperature (20.5**°**C +/-1) as the speed of Drosophila developmental processes, including germband extension are temperature dependent (Butler, 2008).

Wildtype embryos survived to larval hatching post acquisition. *twi* homozygous embryos were identified by reduced mesoderm contractility and failed invagination and confirmed by their cuticle phenotype post acquisition.

### Movie Tracking

First, in order to track at comparable apico-basal positions in all cells regardless of embryo curvature, acquired z-stacks were converted into stacks of curved quasi-two-dimensional representations, following the curved surface of the embryo at each timepoint. Next, automated segmentation, with manual intervention to increase tracking accuracy was performed as previously (Tetley et al., 2016). We segmented cell contours in the Gap43Cherry channel at the level of adherens junctions. During germband extension we used apical myosin signal as a guide to ensure we were segmenting at the correct plane. Note that prior to germband extension apical myosin is very weak and apical adherens junctions are slightly more basal, based on the ruffled appearance of cell membranes more apically and imaging data of cadherinGFP in other datasets form our laboratory (data not shown). Therefore in earlier time frames we used images from up to 4µm deeper into the embryo to accurately segment cell contours.

Any inaccurately tracked cells were removed prior to analysis based on cell velocity relative to neighbours, cell area, cell area change and number of frames over which the cell could be tracked.

The ventral midline (middle of the presumptive mesoderm in *twist)* was followed over time and used to define the orientation of the AP axis, and the DV axis was set perpendicular to this. ‘Distance from mesectoderm’ is defined as distance from the first row of centroids of ectodermal cells at 30 minutes of germband extension.

*Note on number of cells in movies (see Fig. 1-S3)*

In wildtype movies initially the mesoderm takes up a large part of the field of view leaving ectoderm only visible on one side of the embryos at the start of the movies. As the mesoderm invaginates the ectoderm on the other side of the embryo comes into view, and the number of ectodermal cells visible continues to increase as mesoderm invagination completes and ectodermal converge towards the midline during germband extension. Note that the ectoderm that is visible often moves slightly dorsalwards and out of the field of view prior to mesoderm invagination, accounting for the reduction in cell numbers between -15 and 0 minutes of germband extension.

In *twist* movies cell numbers vary less throughout the course of the movie because the mesoderm primordia is narrower in *twist* than in wildtype so takes up less space initially, but does not invaginate successfully so still takes up part of the field of view throughout germband extension. Precise cell numbers vary slightly between movies due to differences in exact DV location of mounting of the embryo and slight variations in the number of cells successfully tracked.

### Movie Synchronisation

Movies of the same genotype were synchronised together using a threshold of 0.01pp/min AP-projected tissue strain rate. This non-zero threshold was chosen due to some minor fluctuations in this strain rate around this start of GBE. Average AP-projected tissue strain rate curves were then calculated for each genotype, and used to synchronise movies so that average AP-projected tissue strain rate was 0 pp/min at time = 0 mins GBE.

For the purposes of checking synchronisation, the appearance of apical myosin in the ectoderm was assessed by eye as signal discernible above background in raw movies. Cytokinesis timing was assessed through by-eye observation of cytokinesis rings.

### Exclusion of mesoderm and mesoectodermal cells from analysis

Cell types were initially broadly defined using rules based on their proximity to the midline at a given timepoint, then were refined by manually selecting cells of incorrect cell-types based on their patterns of cell division within the time period of analysis and redefining them where necessary. Note that in wildtype the width of the mesoderm is approximately 18 cells whilst in the *twist* mutants the width of the mesoderm is reduced to approximately 8-10 cells, and this was taken into account when defining initial midline proximity rules (Leptin and Grunewald, 1990).

For cell division timing we used comparison to fixed embryos with mesectodermal (sim HCR) and dividing cells (anti phospho-H3) marked to ensure accurate cell type identification throughout the analysed time period (Fig1-S1 and Fig1-S2, see HCR methods below). The pattern of cell division was of particular use for the *twist* movies because the boundary between ectoderm and ‘mesectodermal’ cells is not as straight as in wildtype.

### Exclusion of cells stretching in AP due influence of cephalic furrow formation and posterior midgut invagination

For quantitative comparison of AP cell elongation rates between wildtype and twist (Figure 5C) we excluded cells undergoing substantial stretching due to cephalic furrow formation and posterior midgut invagination, so that only AP cell elongation in the central part of the germband was compared. Because of slight variations in the AP positioning of embryos in the field of view we used set individual rules for each movie defining the central region between fixed fixed AP positions within the field of view. We used spatiotemporal plots (AP cell elongation versus AP position and Time of GBE) of individual movies (not shown) as a guide to do this. Note also that movies are not registered precisely along the AP axis for the spatiotemporal plots shown in Figures 5B and 5D due to lack of suitable registration points in our data.

## Movie analysis

### Analysis of cell and cell junction behaviour

Cell shape strain rates were calculated using minimisation to find the best mapping of cell shapes in adjacent pairs of movie frames. Then tissue, cell shape change cell intercalation and cell area strain rates were calculated as previously (Butler et al., 2009; Lye et al., 2015). Axial shape elongation was calculated as previously (Butler et al., 2009; Finegan et al., 2019). Straight vertex to vertex interface lengths and orientations were calculated as previously (Finegan et al., 2019). Rates of change are calculated from 2 time-points (1 minute) before to 2 time-points after the time-point in question.

Neighbour exchange and productive neighbour exchange analysis were calculated as previously (Finegan et al., 2019) except with a modified rule for whether a T1 was productive (extending in AP) or counter-productive (extending in DV). Previously, T1s ware defined as productive when the angle of the centroid-centroid line between the pair of cells that make a new contact was greater than 45 degrees from AP. Here, rather than discrete, we made each T1 contribution continuous, depending on the angle of the centroid-centroid line because we observed that the orientation of *twist* T1s were less strictly aligned with embryonic axes than in the wild-type. T1 contributions ranged continuously from 1 for a centroid-centroid line aligned with DV, through 0 at 45 degrees, to -1 if aligned with AP. Both neighbour exchange measures were expressed as a proportion of the total number of DV-oriented interfaces and as a rate per minute so that they were directly comparable to our intercalation strain rates in units of proportion per minute.

### Myosin quantification

Fluorescence quantification was performed on raw data from the Myosin channel, that had been corrected for artefactual domed Myosin intensity across the field of view as previously (Tetley et al., 2016) and normalised as follows. Background signal was subtracted by setting pixels of intensities up to 5 percentile set to zero for each timepoint. Intensities varied slightly between experiments due to differences in laser intensity and therefore histograms of pixel intensities were stretched: the 98.5 percentile of the data at 30 minutes GBE was set to a greyscale value of 200 with lower value pixels scaled by the same factor and leaving the top 55 values for a tail of high value pixels. We choose to stretch histograms using a focal timepoint, rather than on a per timepoint basis, to preserve the observed increase in Myosin intensity over time.

Tracking of the Gap43 channel was used to define positions of cell interfaces. Because channels were imaged sequentially, images are slightly out of register with each other due to the time elapsed between capturing each channel. Therefore channel registration was corrected post-acquisitionally in order that information on the position of interfaces in the Gap43 channel could be used to locate them in the Myosin channel. Therefore the local flow of cell centroids between successive pairs of time frames in the Gap43 channel is used to give each interface/vertex pixel a predicted flow between frames.

A fraction of this flow is applied, equal to the Myosin II to Gap43 channel time offset, divided by the frame interval. Because cells deform as well as flow, the focal cell’s cell shape strain rate is also applied, in the same fractional manner as above.

We quantified Myosin II maximum intensity projections of 5 z-planes from quasi-two-dimensional stacks extracted prior to tracking (see Movie Tracking) from 2µm above to 2 µm below the level of the adherens junctions during GBE. These thicknesses and projection depths were chosen to capture the majority of junctional-associated myosin whilst minimising contamination with non-junctional signal.

Cell interfaces were defined as a 1-pixel thick line surrounding each cell, with values for shared interfaces averaged between these two 1-pixel thick lines. Because cell vertices are often enriched in Myosin, vertex pixels (within a radius of 2 pixels of the vertex point) were removed for the quantification.

We then calculated the bipolarity (planar cell polarity, with 2 peaks approximately at opposite cell edges) of each cells average interface myosin by fourier analysis, extracting the average amplitude of two Myosin peaks when plotting the interface fluorescence intensity around the cell perimeter. Prior to fourier analysis, elongated cells were unstretched to eliminate bias introduced by unequal interface lengths. The phase of the period 2 component of the fourier series gives the direction of the bipolarity. The bipolarity is presented as both unprojected values and projected to the embryonic anterior-posterior axis. Rates of change are calculated from 2 timepoints before to 2 timepoints after the timepoint in question.

### Statistics and graphical representations

Ribbon plots were generated in R with 30 second bins and profiles smoothed over 3 bins for presentation only (significance tests were calculated on unsmoothed data). We used a mixed effects model to test for differences between genotypes with a p-value of 0.01 as previously (Finegan et al., 2019). Significant differences are highlighted in grey on plots. Ribbons show standard error between genotypes. Spatiotemporal plots were generated in otracks(Blanchard et al., 2009). Other plots (neighbour exchanges, synchronisation plots) were generated in Prism with 1 minutes bins.

## ACKNOWLEDGEMENTS

We acknowledge Bruno Monier, Stefano De Renzis and Bloomington Drosophila Stock Center (https://bdsc.indiana.edu/) for Drosophila strains. We thank all members of Bénédicte Sanson’s research group for useful discussions. This work was supported by Wellcome Trust Investigator Awards to B.S. (099234/Z/12/Z and 207553/Z/17/Z).

## SUPPLEMENTARY FIGURE AND MOVIE LEGENDS

**Figure 1-S1:**
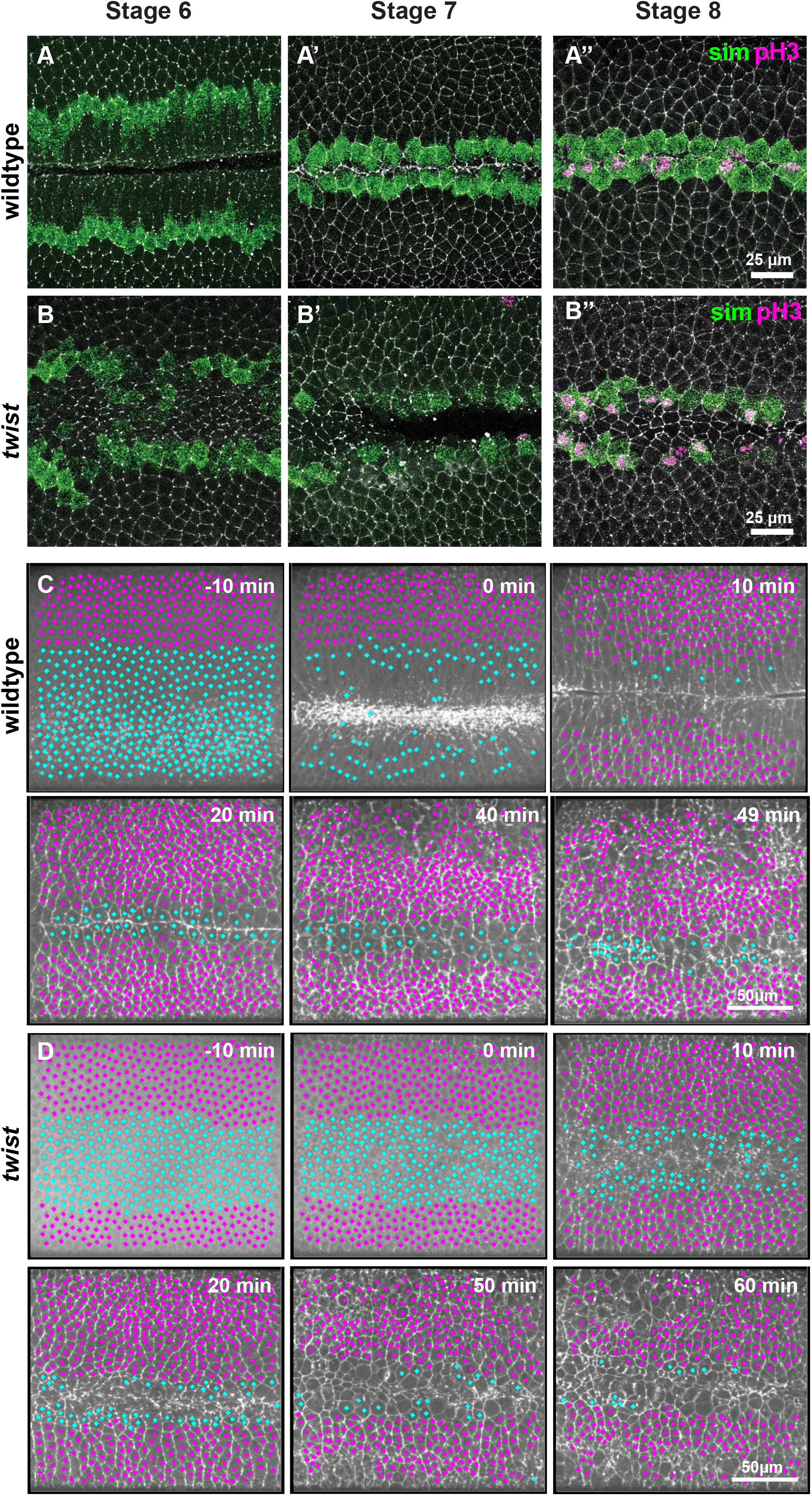
Defining cell types in wildtype and *twist* mutant movies. (A, B) Representative images of HCR of *single-minded* (*sim*, green, mesectoderm) with antibody staining of phosphoHistone3 (pH3, magenta, dividing cells) in stage 6 to 8 fixed embryos for wildtype (A) and *twist* (B), used to inform definition of cell types in movies (examples in C, D). (C, D) Examples of defining cell types in wildtype (C) and *twist* (D) movies at representative timepoints throughout the course of the movie (mesoderm/mesectoderm, cyan; ectodermal germband, magenta), overlayed on Myosin II channel (maximum intensity projection). Unmarked cells are poorly tracked and excluded from the analysis.

**Figure 1-S2:**
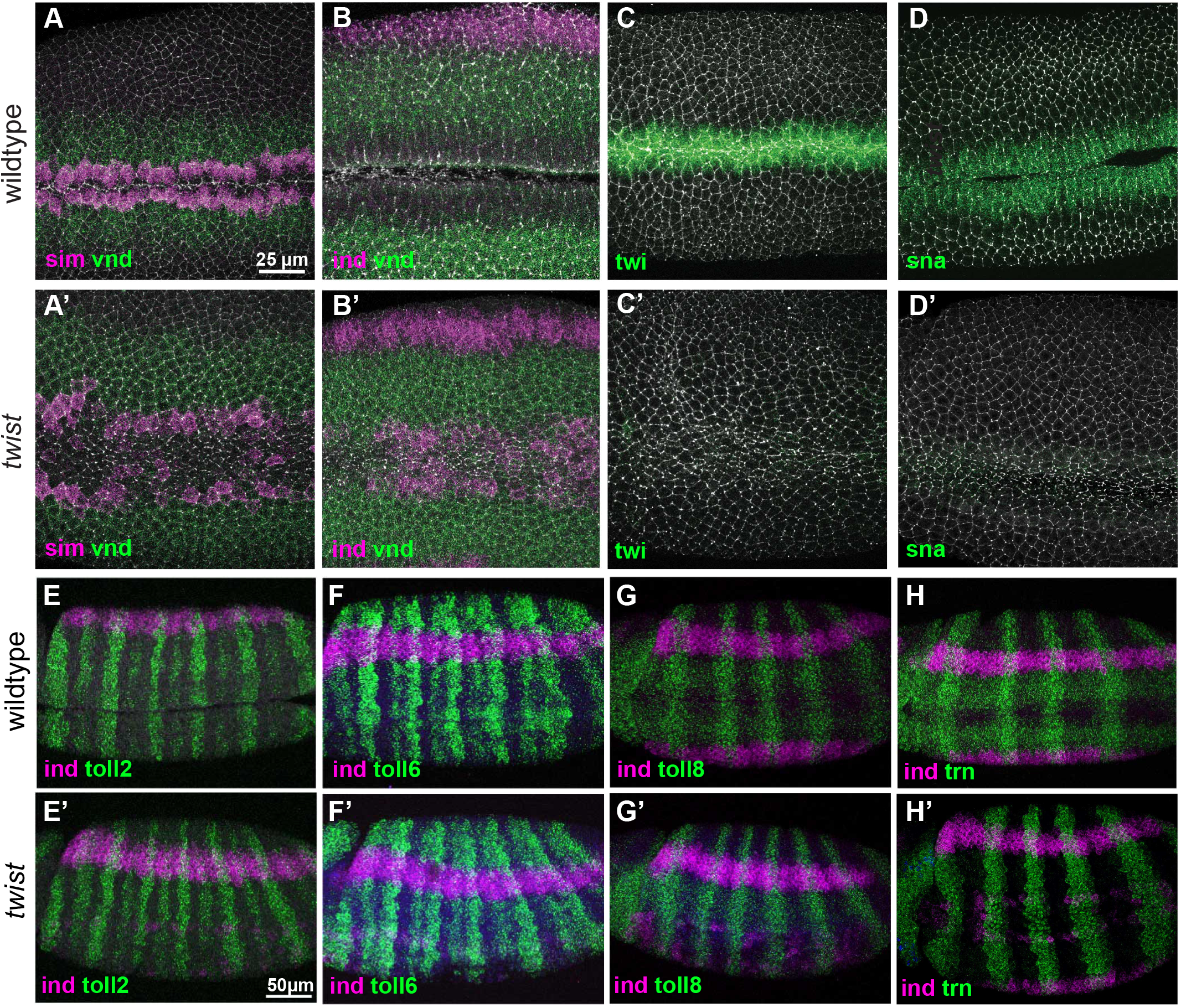
Expression of DV patterning genes and LRR receptors in wildtype and *twist* mutants. (A-D) HCR of DV patterning genes in wildtype and *twist* mutant (A’-D’) stage 7 embryos: (A and A’) *sim* (magenta) and <u>v*nd*</u> (green); (B and B’) *ind* (magenta) and *vnd* (green); (C and C’) *twist* (green), which is missing in *twist* mutants (D and D’) *snail* (green) which is reduced and patchy in *twist* mutants as expected. HCR of LRR receptors in wildtype (E-H) and *twist* mutant (E’-H’) stage 7 embryos (green): (E and E’) Expression of *Toll-2*; (F and F’) Expression of *Toll-6*; (G and G’) Expression of *Toll-8*; (H and H’) Expression of *tartan* (*trn*). Maximum intensity projections of approximately 20µm image stacks. (A-D’) are also marked with anti-phosphoTyrosine (pTyr, white, cell membranes). (E-H’) are also stained for *ind* expression (HCR, magenta) and marked with phosphoHistone3 (pH3, blue, dividing cells).

**Figure 1-S3:**
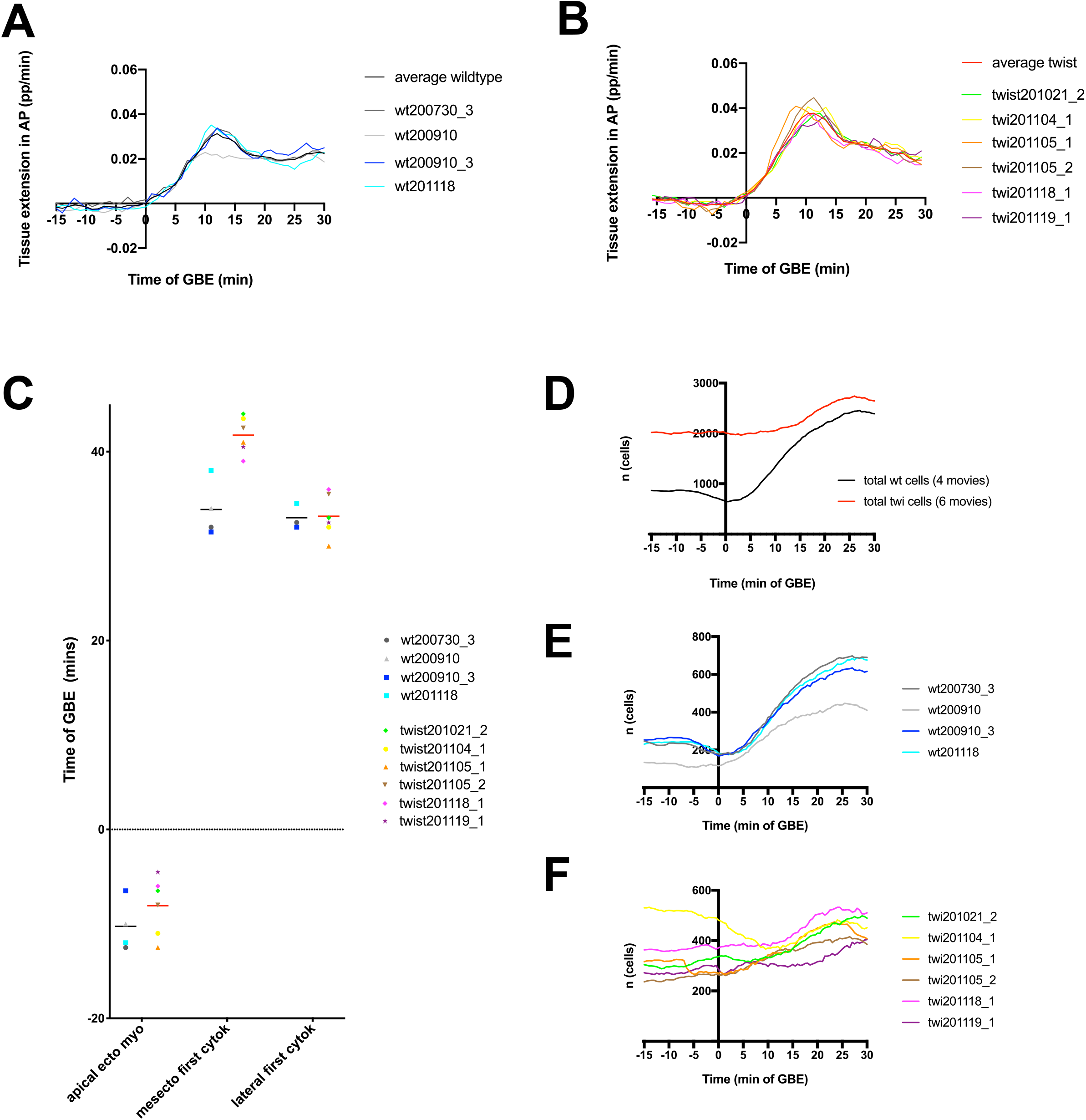
Numbers of cells analysed in wildtype and *twist* mutant movies. (A, B) Synchronisation of movies to start of germband extension using tissue strain rates (proportion/minute), projected along AP axis in wildtype (A) and *twist* (B). (C) Manual check of synchronisation using key developmental events visible in our field of view. Note that the movies are longer than the -15 to 30 minutes of germband extension analysed and vary slightly in length. Note data is missing for a first lateral cytokinesis of *wt200910* it is due to the movie finishing too early for this to be assessed. Lines show average time of developmental events (black, wildtype; red, *twist*). (D-F) Number of ectodermal cells analysed in wildtype and *twist* mutant movies over time. (D) Total number of cells analysed in all movies; (E) number of ectodermal cells analysed per wildtype movie over time and (F) number of ectodermal cells analysed per *twist* movie over time.

**Figure 2-S1:**
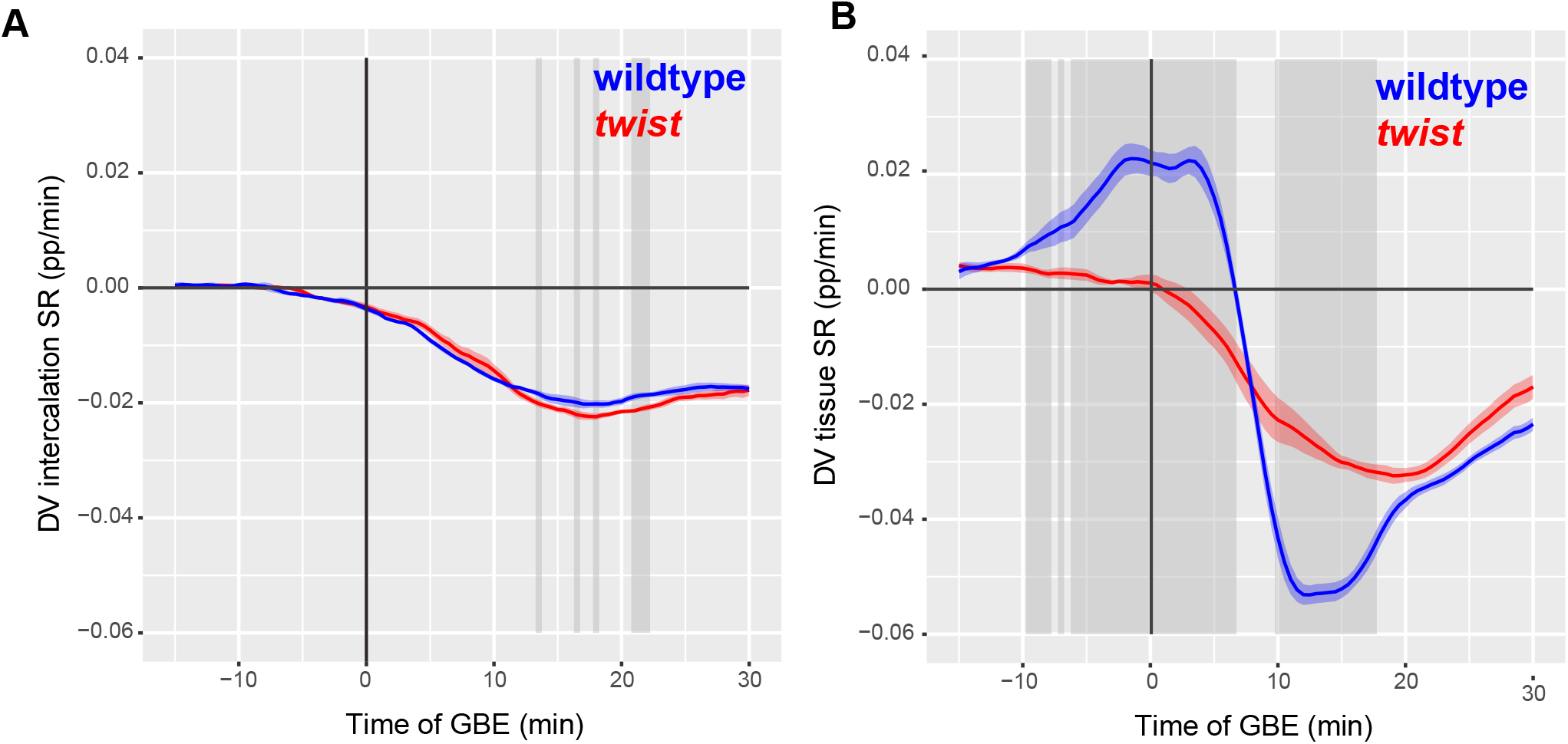
Germband cells do not slip past each other in response to pull from mesoderm invagination. (A) DV-projected intercalation change strain rate (proportion/minute) of all tracked ectodermal cells, summarised for wildtype (blue) and *twist* (red) over time of GBE. (B) DV-projected tissue shape change strain rate of all tracked ectodermal cells, summarised for wildtype (blue) and *twist* (red) over time of GBE. Note that Intercalation Strain Rate is calculated as Tissue Strain Rate minus Cell Shape Strain Rate (Blanchard et al., 2009). Therefore, DV Tissue Strain Rate (B) is very similar to DV Cell Shape Strain Rate (Fig. 2C) before the start of germband extension as DV Cell Intercalation Rates are close to zero between -15 and 0 minutes of GBE.

**Figure 3-S1:**
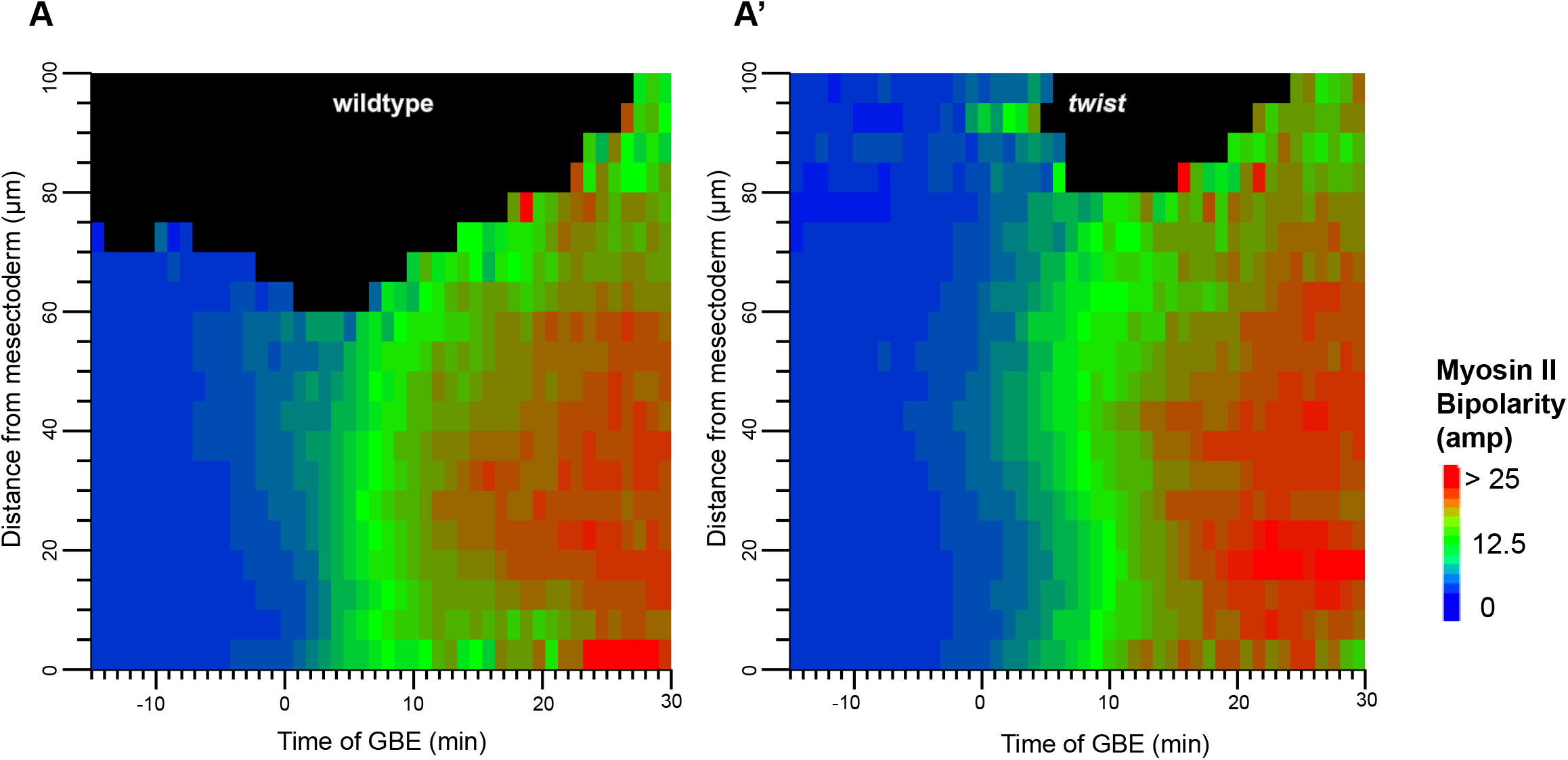
Spatiotemporal plots of Myosin II bipolarity in wildtype and *twist* mutants. Unprojected Myosin bipolarity of all tracked ectodermal cells, summarised for wildtype (A) and *twist* (A’) against time of GBE and distance from mesectoderm (µm) (see Methods).

**Figure 5-S1:**
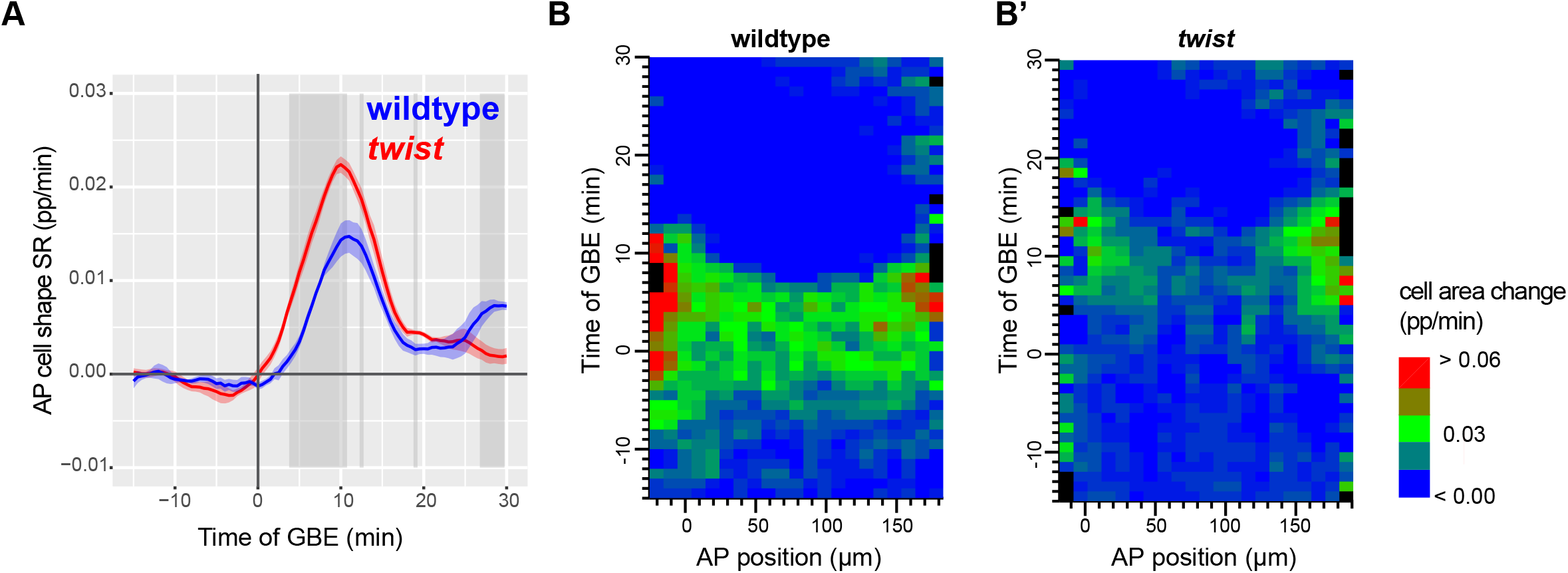
AP cell elongation and cell area change in wildtype and *twist* mutants. AP cell shape change strain rate plotted time of GBE summarising data from the full imaged region of 4 wildtype (left) and 6 *twist* (right) movies. (B, B’) Spatiotemporal plots of cell area change strain rate plotted against anterior-posterior position and time of GBE extension summarising 4 wildtype (B) and 6 *twist* (B’) movies.

**Figure 6-S1:**
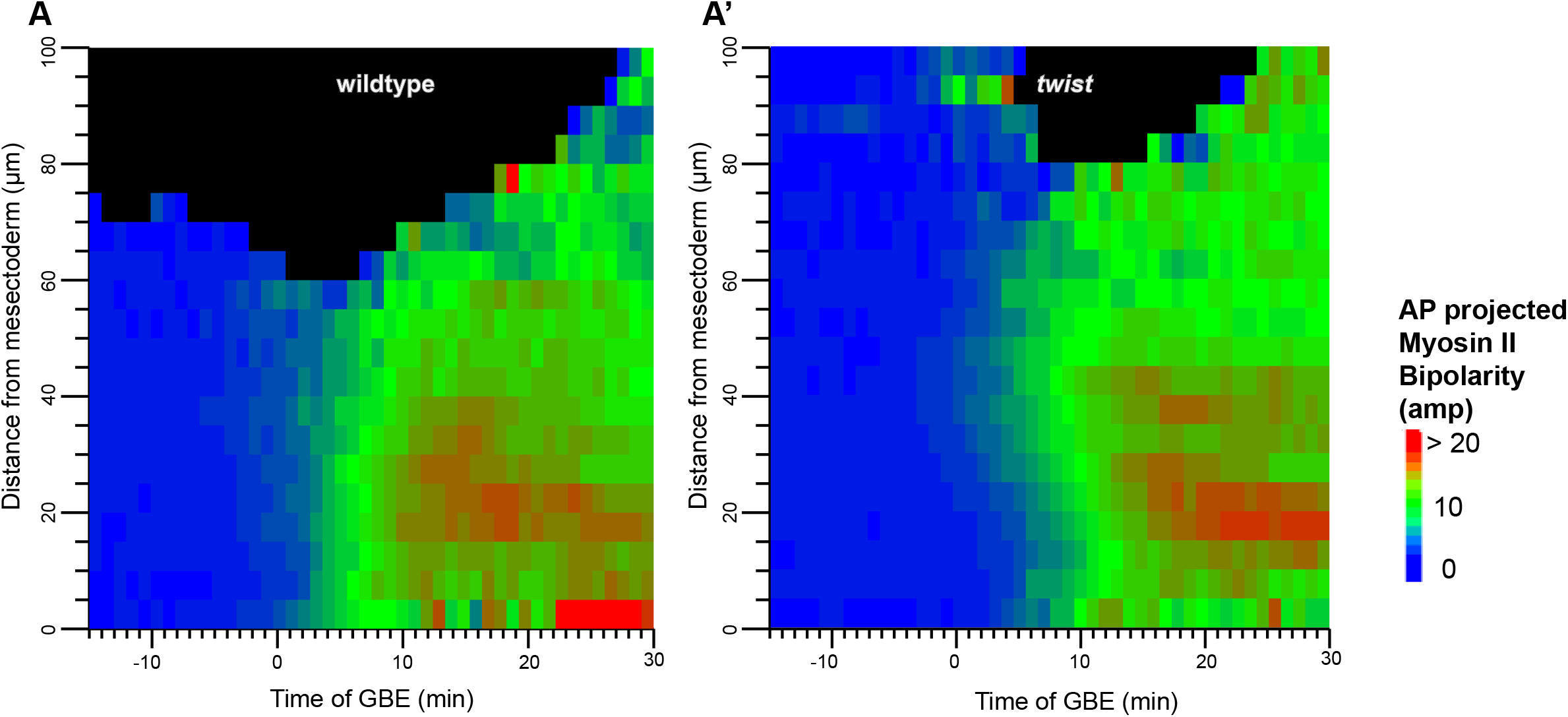
Spatiotemporal plots of projected Myosin II bipolarity in wildtype and *twist* mutants. (A, A’) Myosin bipolarity projected along AP of all tracked ectodermal cells, summarised for wildtype (A) and *twist* (A’) against time of germband extension and distance from mesectoderm (µm) (see Methods).

**Movie 1: Example of cell tracking in wildtype (prior to exclusion of mesoderm/mesectoderm)**

Example wildtype (wt201118) movie from -15 to 30 minutes of germband extension, showing manually corrected tracked cells with additional quality control rules applied to remove inaccurately tracked cells and incomplete cells at the edge of the embryo (see Methods). Cell contours (magenta) and centroids (white) overlayed on Gap43Cherry signal from the apical adherens junction level of the embryo, which was extracted by ‘blanketing’ of image stacks (see Methods; *Movie Tracking*).

**Movie 2: Example of cell tracking in *twist* (prior to exclusion of presumptive mesoderm/mesectoderm)**

Example twist (twi201118_1) movie from -15 to 30 minutes of germband extension, showing manually corrected tracked cells with additional quality control rules applied to remove inaccurately tracked cells and incomplete cells at the edge of the embryo (see Methods).Cell contours (magenta) and centroids (white) overlayed on Gap43Cherry signal from the apical adherens junction level of the embryo, which was extracted by ‘blanketing’ of image stacks (see Methods; *Movie Tracking*).

